# Gene expression signature of castrate resistant prostate cancer

**DOI:** 10.1101/2022.03.16.484397

**Authors:** JM Dixcy Jaba Sheeba, Shraddha Hegde, Ninad Tamboli, Namratha Nadig, Ramaiah Keshavamurthy, Prathibha Ranganathan

## Abstract

Prostate gland is a highly androgen dependent gland and hence the first line of treatment for metastatic prostate cancer happens to be androgen ablation. This is achieved by multiple non-surgical methods. However, most of these cancers although respond well initially, become resistant to androgen ablation sooner or later. These cancers then become extremely aggressive and difficult to treat, thereby drastically affect the patient prognosis. The purpose of this project was to identify a gene expression signature for castrate resistant prostate cancer which may aid in identification of mechanisms responsible for castrate resistance. For this purpose, we have collected patient samples belonging to a. Control group; b. Castrate Sensitive group and c. Castrate resistant group. Gene expression profiling has been done on these samples using RNA-seq and several differentially expressed genes identified between the castrate sensitive and resistant groups. We have also identified some genes which are expressed in the castrate resistant group alone, which is of interest since these may have an implication in evolution of castrate resistance and also prognosis.

We have compared this with data from The Cancer Genome Atlas (TCGA). Using criteria such as overall survival, disease free survival, progression free survival and biochemical recurrence, we have identified genes which may have relevance in progression to castrate resistance and in prognosis. Functional annotation of these genes may give an insight into the mechanism of development of castrate resistance.

## Introduction

Prostate cancer is the second most common cancer in men and the second leading cause of cancer related deaths in the United States [1]. Prostate cancer accounts for 3.8% of all cancer related deaths world-wide [2]. Although there are many cell types in the prostate gland which can give rise to cancer, the most common one is adenocarcinoma. Many lines of treatment approaches are followed to treat the disease such as surgery, radiotherapy, chemotherapy, hormonal therapy, depending on the stage of disease. There are also other kinds of therapy such as immunotherapy, cryo-surgery, photon beam therapy, which are used in certain cases. Given that prostate is highly dependent on androgens for its normal development and functioning, androgen deprivation therapy is a choice therapeutic strategy, particularly for metastatic prostate cancer. This is achieved by a. Blocking androgen production by the testicles (using surgical castration or estrogens or LHRH analogs); b. Blocking androgen synthesis throughout the body (inhibition of CYP17, an enzyme required for synthesis of testosterone from cholesterol) or c. By blocking the action of androgens using receptor blockers such as flutamide, bicalutamide etc [1],(Reviewed in [3–5]). Although Androgen Deprivation Therapy (ADT) gives good results in the beginning, many cases develop resistance to this therapy. This is usually manifested as high levels of Prostate Specific Antigen (PSA) despite androgen deprivation [6, 7]. Once the cancer reaches this stage, it is difficult to manage and has poor prognosis. Several mechanisms have been proposed for development of castrate resistance (reviewed in [8]). This includes amplification of the AR (androgen receptor) gene, mutations in AR, splice variants, mutations in the co-activators and repressors, all leading to aberrant AR signaling. There have been several studies on prostate cancer including castrate resistant prostate cancer which have identified genetic aberrations which may be indicators of progression and prognosis of prostate cancer. Whole Genome Sequencing studies have shown that alterations in RB1 and β-catenin genes are associated with poor prognosis as well as resistance to enzalutamide treatment [9]. Another study on patient tumor DNA has revealed that somatic mutations in the Wnt pathway genes such as CTNNB1, APC or RNF43, that can lead to activation of the pathway correlate with poor prognosis and resistance to ADT [10]. Mutation studies on cell free tumor DNA from plasma of prostate cancer patients has revealed mutations in PI3K, CTNNB1 correlate with poor prognosis. Besides, the study has also shown certain mutations in the AR gene such as L702H and T878A enriched in CRPC [11]. There are also many studies which have profiled gene expression in prostate cancer [9, 12, 13]. However, despite the fact that any variations in AR or AR signaling would eventually affect gene expression, there are very few studies comparing the gene expression profiles of castrate sensitive and resistant prostate cancers. Moreover, when we consider the Indian scenario, although we see steady increase in the prostate cancer incidence, there are no published profiles representing this population. Our study was designed at addressing these two lacunae, i.e to get a comparative gene expression signature between castrate sensitive and resistant cancers and also a gene expression profile of Indian population.

## Materials and Methods

### Patient specimen

All patient specimen were collected from the department of Urology, Institute of Nephro-Urology, Bengaluru. The samples were collected either during TRUS/TURP procedures. The control group comprised of patients who had been diagnosed of Benign Prostatic Hyperplasia and confirmed by pathologist to be non-malignant. The cancer samples included in this study were from the patients who had been diagnosed as metastatic carcinoma prostate and on Androgen Deprivation Therapy(ADT). ADT includes surgical castration in the form of bilateral orchiectomy or medical castration with luteinizing hormone releasing hormone (LHRH) agonist/antagonist or Combined Androgen Blockade (CAB). These patients were followed up on a regular basis with clinical examination and serum PSA levels. If patients’ PSA levels were on rising trend then they were called for follow up according to EAU guidelines [14] for diagnosing castrate resistance.

These patients were then classified as Castrate sensitive (CaS) and Castrate resistant (CaR) according to the EAU guidelines [14]. Patients who maintained nadir PSA levels on follow up after ADT were classified as Castrate Sensitive. The patients who had castrate serum levels of testosterone < 10 ng/dl with 3 consecutive rises of PSA, 1 week apart resulting in two 10 % increases over the nadir were classified as Castrate Resistant.

In our study we have used 31 controls, 23 CaS and 4 CaR samples, all collected between 2018-2021.

### RNA isolation

All tissue samples were collected in RNA later (Ambion Inc, USA). Prior to RNA isolation, the tissues were removed from RNA later, rinsed with PBS and homogenized in RLT buffer (QIAGEN GmbH) using a hand held homogenizer. RNA isolation was done using Qiagen RNAeasy Mini Kit according to manufacturer’s instructions (QIAGEN GmbH). The RNA was quantified using Nano-drop (Thermo Scientific, USA) and 2-3ug RNA was used for RNA-seq analysis.

### RNA-sequencing

The RNA sequencing was outsourced to Sandor Life Sciences Pvt Ltd, Hyderabad and Institute of Bioinformatics and Applied Biotechnology, Bengaluru. After initial QC analysis, samples which qualified the criteria were processed further for library preparation. The sequencing was done on Illumina Platform with 20-25 million paired end reads.

### Analysis of the RNA-seq data

The analysis of the RNA-seq data was done with the help of Shodhaka Life Sciences, Bengaluru.

In brief, around 260 Giga bytes of RNA-sequencing data from castrate-resistant, castrate-sensitive and control samples was assessed for quality. High quality reads obtained after a thorough filtering process were aligned against the latest hg38 human reference genome and transcriptome and used for quantifying genes and transcripts. Differential expression analysis was performed and significantly differentially expressed genes with a minimum log2 fold of 1.2 and p-value <0.05 were obtained.

### Identification of castrate-resistant or castrate-sensitive genes in prostate cancer samples

Using the data obtained from our RNA-Seq analysis, differentially expressed genes in CaR group with respect to BPH were compared with the differentially expressed genes in CaS group with respect to BPH groups. Genes uniquely present in CaR group in comparison with CaS group were identified (CaR genes). Similarly genes uniquely present in CaS group in comparison with CaR group were identified (CaS genes).

### Retrieval of data from The Cancer Genome Atlas (TCGA)

RNA-seq data of prostate cancer and normal tissues were retrieved from The Cancer Genome Atlas [15] database using Xena Browser [16]. The dataset comprised of 550 samples, in which 498 samples are prostate tumors, and 52 are normal tissues. The average age of the patients was 61 years (range 41–78 years). The gene expression data were generated from Illumina HiSeq 2000 RNA Sequencing platform and were mentioned in terms of log_2_(RPKM+1). Samples were segregated as tumor and normal samples based on the barcode. mRNA expression of each of the differentially expressed (DE) genes in both the groups was analyzed by Welch two-sample t-test in R Software. Genes that are significantly different in both groups were identified.

### Identification of differentially expressed genes relevant in prognosis

For this, we have considered four criteria namely Gleason score, survival, biochemical recurrence and tumor recurrence after initial treatment. The above data were retrieved from TCGA [15] and the differentially expressed genes which correlate with these criteria identified.

### Correlation of expression of DE genes with Gleason Score

Gene expression data of DE genes were merged with clinical data using barcode. Correlation between DE gene expression with the Gleason Score was assessed by Spearman’s rank correlation test in R software. Genes with Spearman correlation coefficient (ρ) > 1 indicates the positive correlation, and ρ < 1 indicates the negative correlation. Genes that are significantly associated with the Gleason score were identified. Positive correlation indicates higher expression in tumor has higher Gleason score.

### Identification of differentially expressed genes across biochemical recurrence

Out of 23530 genes, genes with no expression or low expression values (median = 0) were excluded and 17711 genes were considered for the analysis. Biochemical recurrence data was available for 430 among which 58 patients developed biochemical recurrence. Gene expression data was merged with clinical data using barcode, and samples were dichotomized based on biochemical recurrence. mRNA expression values of 17711 genes corresponding to samples with biochemical recurrence and samples without biochemical recurrence were analyzed by Welch two-sample t-test in R Software. Genes that were significantly different in both the groups were compared with the differentially expressed genes from our data.

### Survival analysis

Survival data of prostate cancer patients were retrieved from the TCGA database. Association of DE genes with Overall survival (OS), progression -free interval (PFI), disease-free interval (DFI) were analyzed using the KM Plotter online tool [17–19]. For each gene, tumor samples were divided into two groups using the median expression value as a cut off and the hazard ratio between the groups was calculated. Genes that are significantly associated with the OS, PFI, or DFI were identified, and Kaplan-Meier analyses for these performed.

#### Identification of differentially expressed genes across tumor recurrence after initial treatment

mRNA expression values of DE genes corresponding to samples with tumor recurrence and samples without tumor after initial treatment were extracted. mRNA expression of each DE gene in both the groups was analyzed by Welch two-sample t-test in R Software. Genes that are significantly different in both groups were identified.

#### Statistical analysis

The difference in the means of the two groups was analyzed by Welch’s two-sample t-test. The correlation between Gleason score and gene expression was analyzed by Spearman’s correlation test. All the statistical analyses were performed using the R statistical package. In all the statistical tests, p < 0.05 was considered significant.

## Results

### Differential expression of genes in castrate sensitive and castrate resistant prostate cancer

Based on the above mentioned criteria, it was found that there were 481 differentially expressed genes in the Castrate sensitive group compared to BPH group. Among these, 261 were over expressed and 220 under expressed in the castrate sensitive prostate cancer (Fig 1, Supp Table 1a). Similarly, there were 446 differentially expressed genes in the Castrate resistant group compared to BPH group. among these, 154 were over expressed and 292 under expressed in the castrate resistant prostate cancer (Fig 1, Supp Table 1b). When compared across groups, we found 82 genes which are common. Among these, there were 32 genes which were commonly increased both is CsPC and CRPC and 44 genes decreased in CsPC and CRPC. 6 genes showed opposite expression in the two groups (Supp Table 1c). The gene lists for each of these groups can be found in the supplementary information.

**Figure 1:**
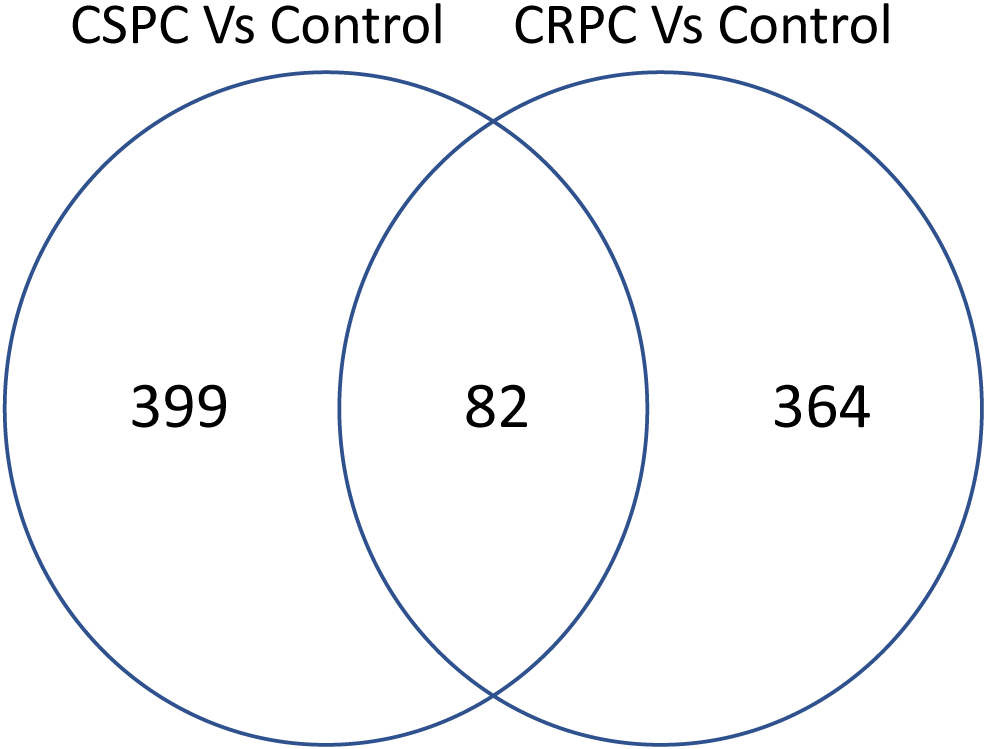
A summary of the differentially expressed genes.

When the gene expression profiles are compared across all the groups, it was seen that 927 genes are differentially expressed in prostate cancer compared to BPH samples. Among these, 415 are over expressed and 512 are under-expressed in prostate cancer samples. 364 genes are uniquely expressed in A.

### Comparsion with data available from public domain

Since our data has been collected over a period of 3 years, we do not have survival data for these samples. Also, our data represents only Indian population. In order to compare it with other populations and also the relevance of this gene expression signature in prognosis, we have compared our data with data from TCGA. TCGA has 20530 genes differentially expressed between normal and cancer with the same criteria as has been considered for our data set. Among the 927 genes which we have found differentially expressed in prostate cancer in our data, 763 genes are also found to be differentially expressed in the TCGA data. However, there are 302 genes which are present in our data for which we do not have data from TCGA (Supp Table 2)

Since we are interested in understanding the prognostic significance of the gene expression signature, we have used only the common genes for all further analyses.

### Identification of genes uniquely expressed in castrate-resistant and castrate sensitive prostate cancer genes

Comparison of differentially expressed genes in CaR group (with respect to BPH) and CaS group (with respect to BPH) showed 364 genes were uniquely expressed in CaR group and 399 genes were uniquely expressed in CaS group. These two gene sets will be represented as CaR genes and CaS genes respectively. 82 genes were differentially expressed in both CaR group and CaS group when compared with BPH group individually Further, log2FC obtained from both these comparisons were analyzed. 6 genes showed opposite direction of regulation and 6 genes showed log2FC difference of greater than 3. To compare our observation with the publicly available data, expression data of CaR genes, CaS genes, and common genes were retrieved from TCGA database. Out of 364 CaR genes, expression data was available for 295 genes, and out of 399 CaS genes, expression data was available for 332 genes (Table 1). We have focused our further analyses on these two sets of genes.

**Table 1:**
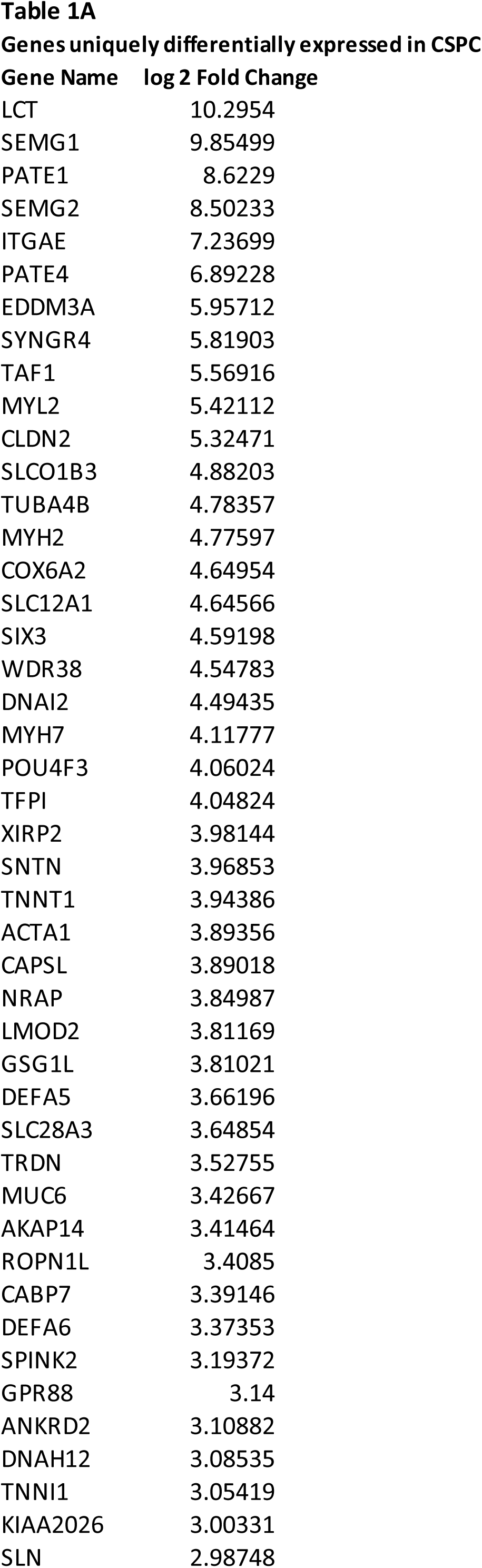
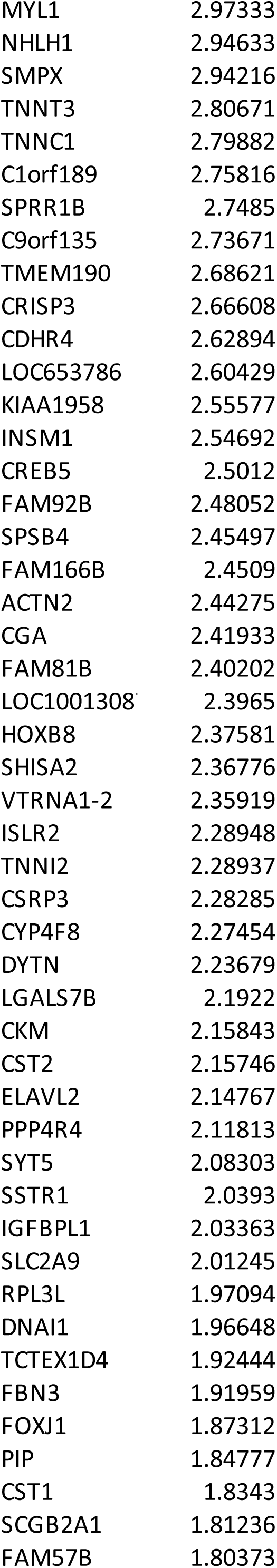
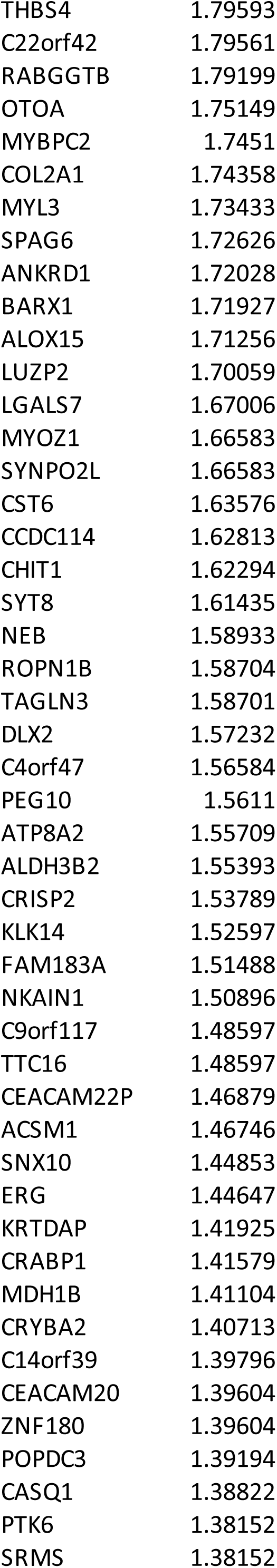
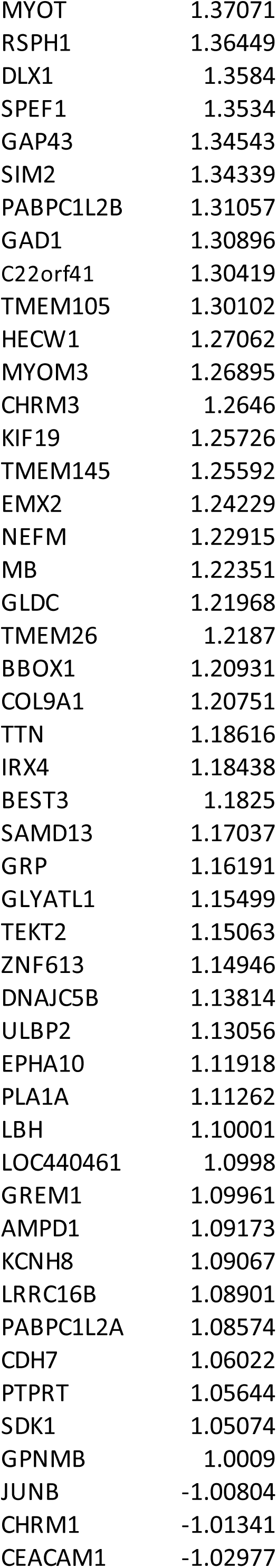
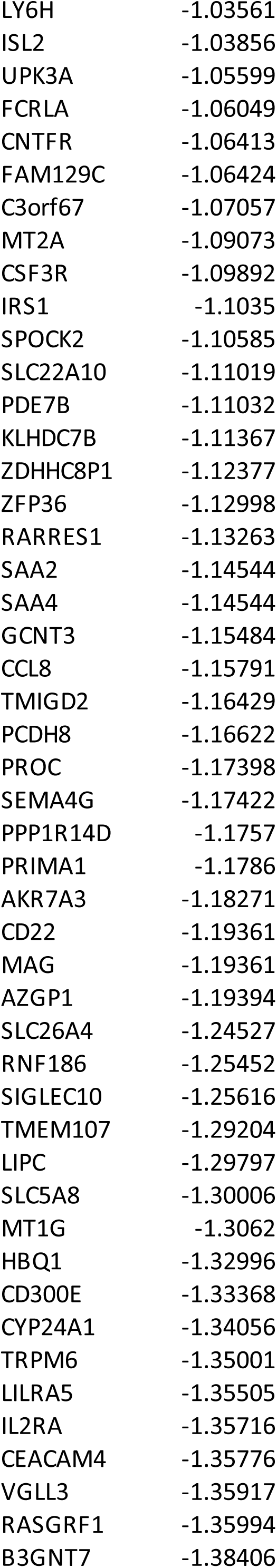
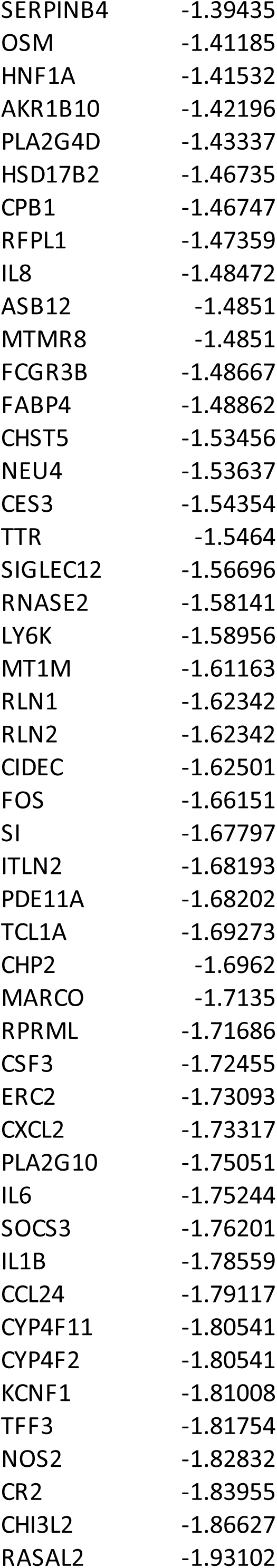
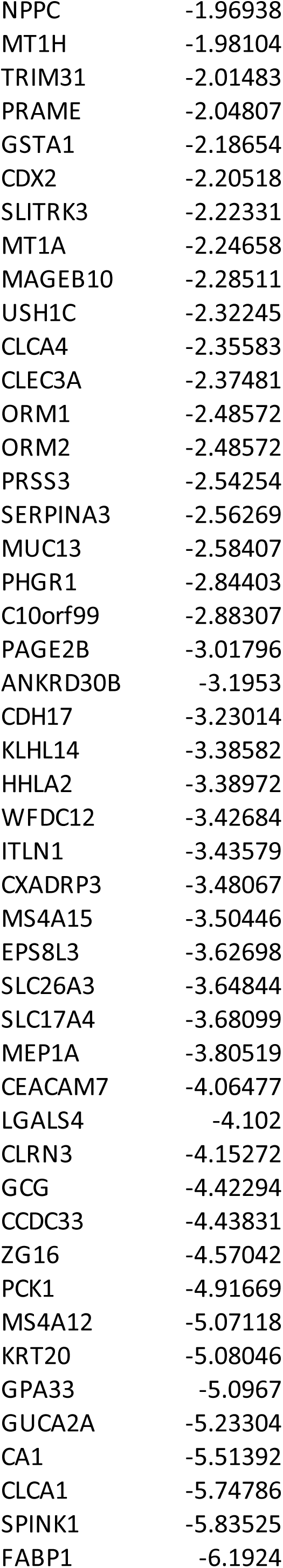
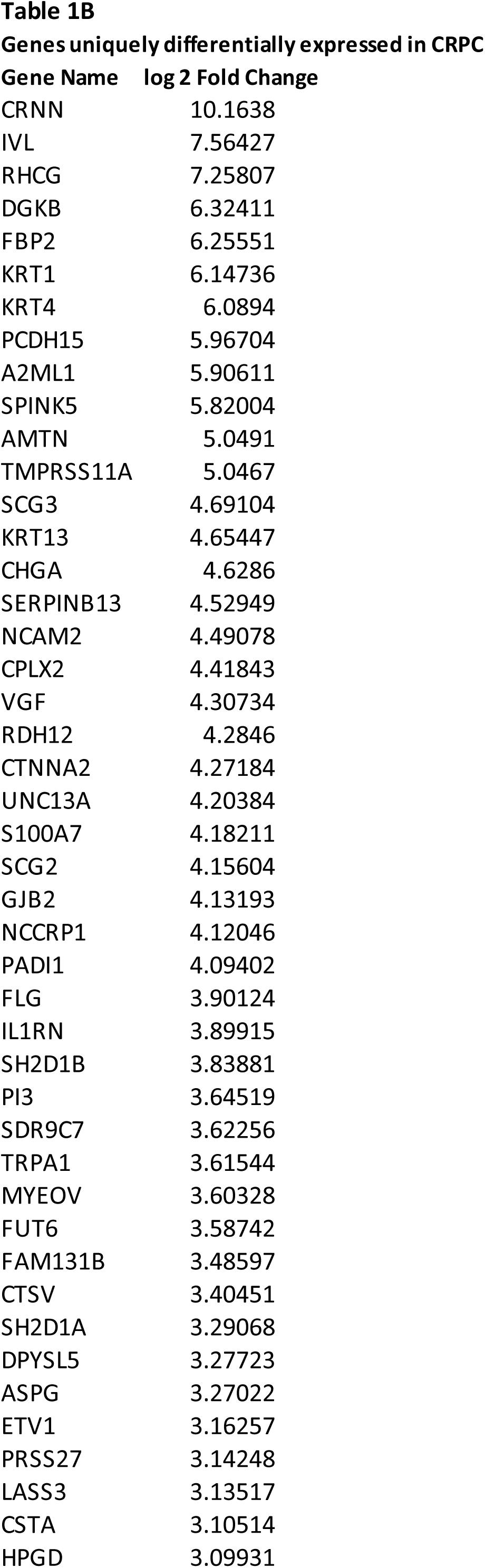
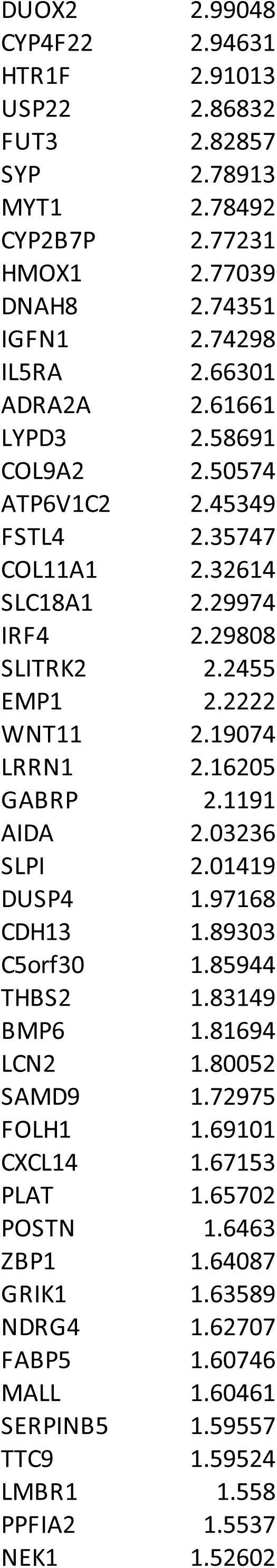
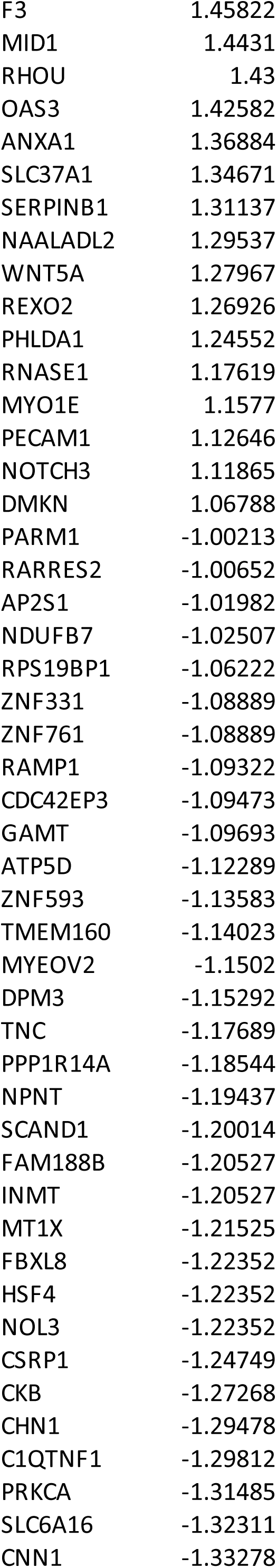
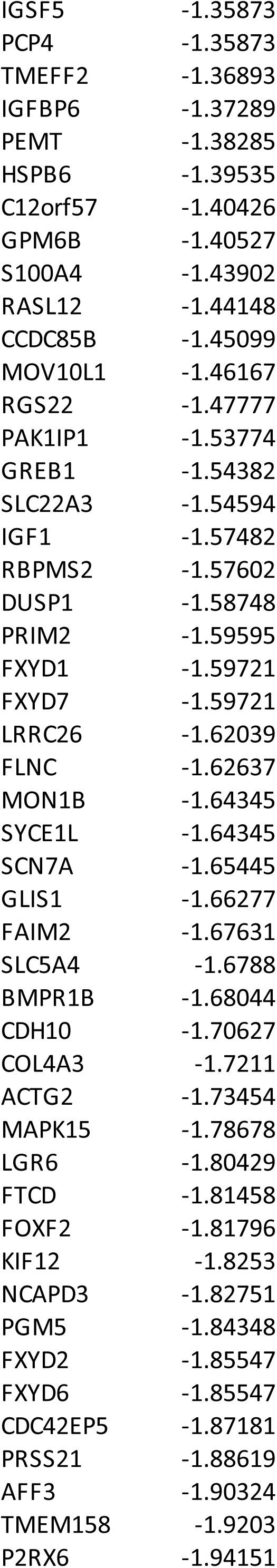
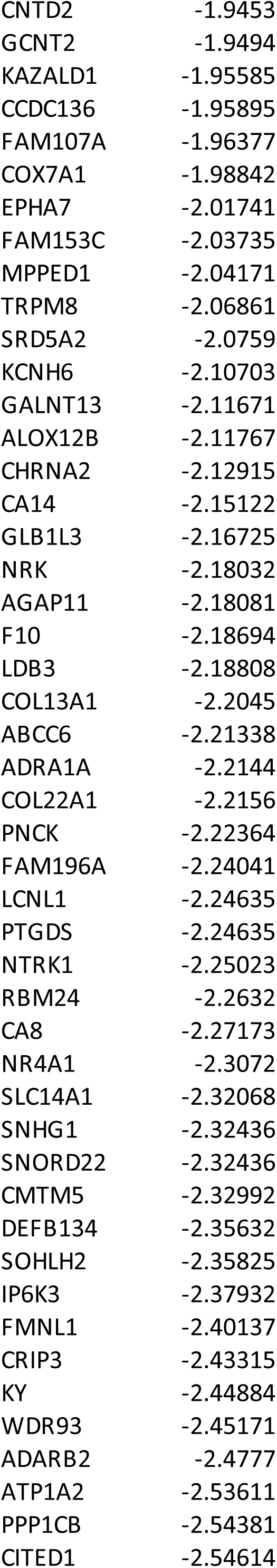
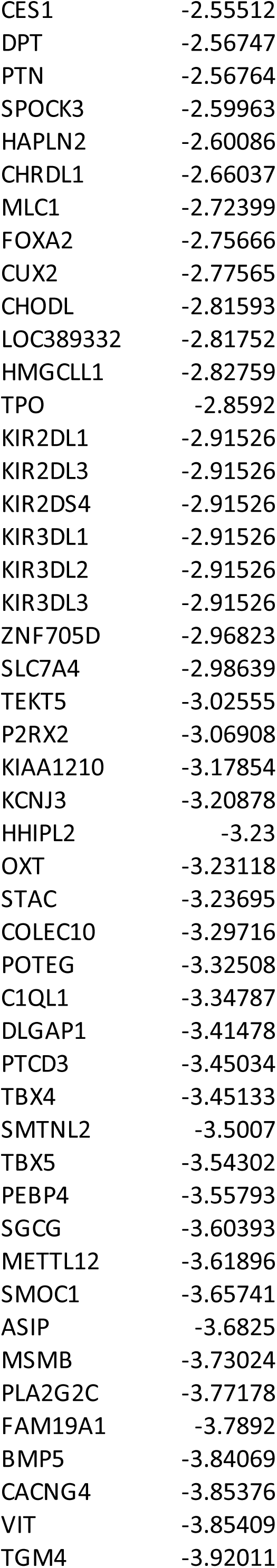
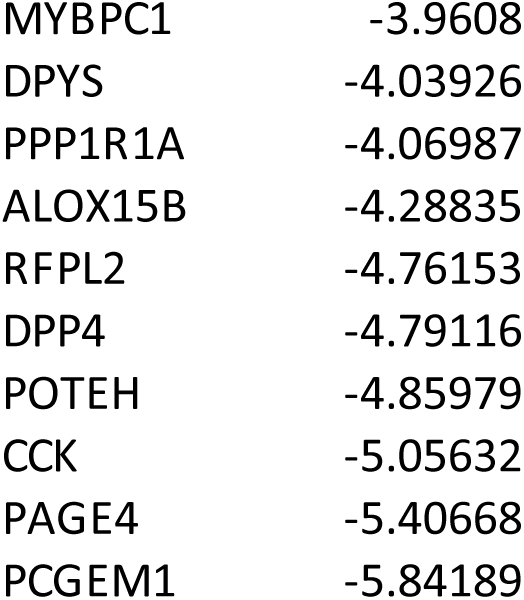
A. List of Castrate Sensitive genes. (332) B. List of Castrate Resistant genes. (295)

### Identification of genes which could be potential biomarkers of prognostic significance

Resistance to androgen deprivation therapy is one of the major challenges in treatment of prostate cancer. In our study we have tried to identify the potential gene expression signature of CRPC. In order to evaluate whether these genes have any value as prognostic markers, we analysed the gene expression data with respect to overall survival, biochemical recurrence, Gleason score and tumor recurrence. These parameters have been retrieved from TCGA.

### Identification of genes associated with biochemical recurrence

**Biochemical recurrence is defined as** an increase in the blood level of PSA in prostate cancer patients after surgery or radiation. This may be an indication that the cancer has relapsed and is also known as PSA failure [1].

In order to identify the genes associated with the BCR, we have used data from TCGA. mRNA expression values of 17711 genes and details about BCR were retrieved from TCGA. in the samples with BCR and samples without BCR were analyzed. 3298 genes were found to be differentially expressed in both groups (p < 0.05). In the samples with BCR, 2006 genes showed higher more expression, and 1292 genes showed lower less expression as compared to samples without BCR

These gene lists were then compared with the differentially expressed gene sets from our data. It was found that 78 CaR genes (27 genes were upregulated and 51 were downregulated in samples with BCR) and 56 CaS genes (18 genes were upregulated, and 38 were downregulated in samples with BCR) Although it appears that more CaR genes are involved in biochemical recurrence, at this point, a direct correlation cannot be made since our data does not have information about biochemical recurrence and also the sample size is small. Table 2 shows a list of genes that may be involved in biochemical recurrence.

**Table 2:**
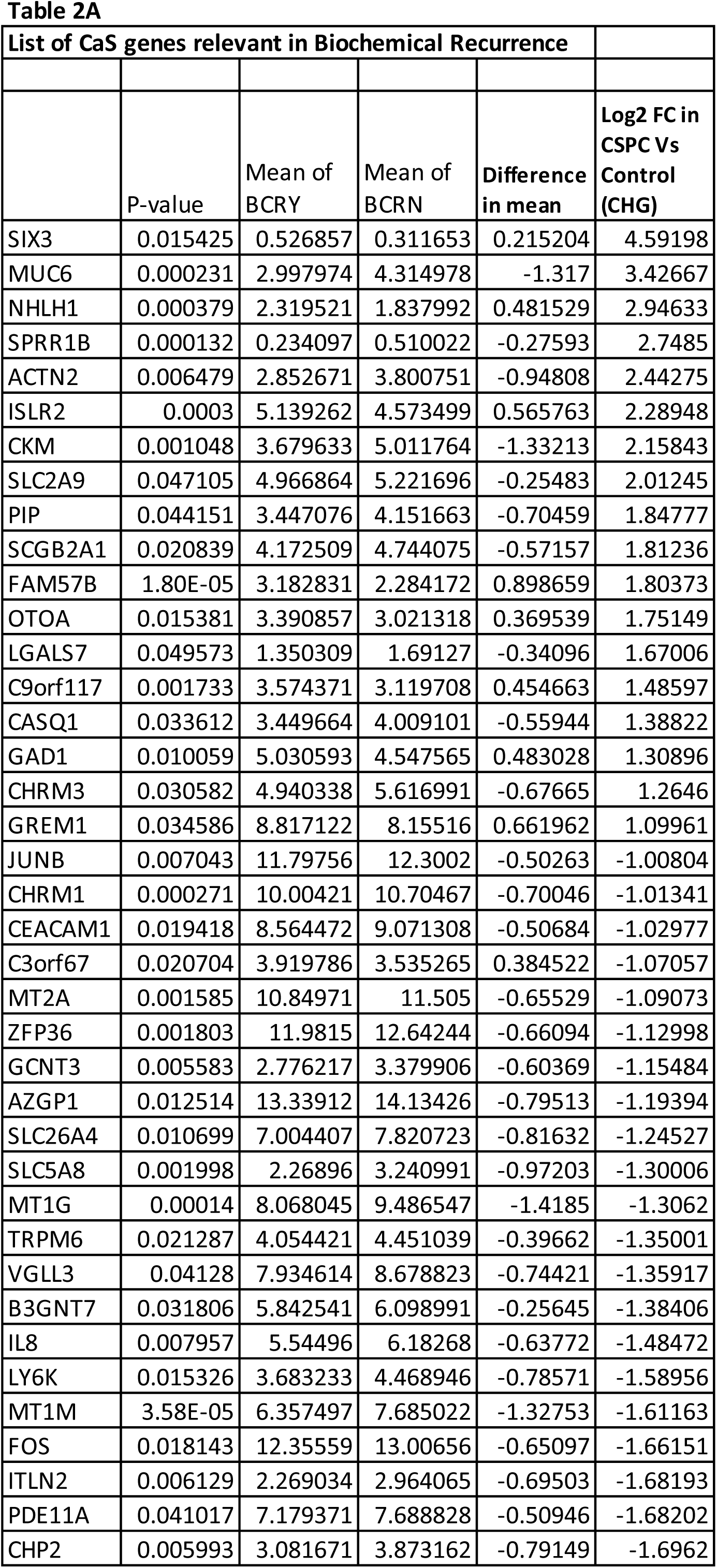
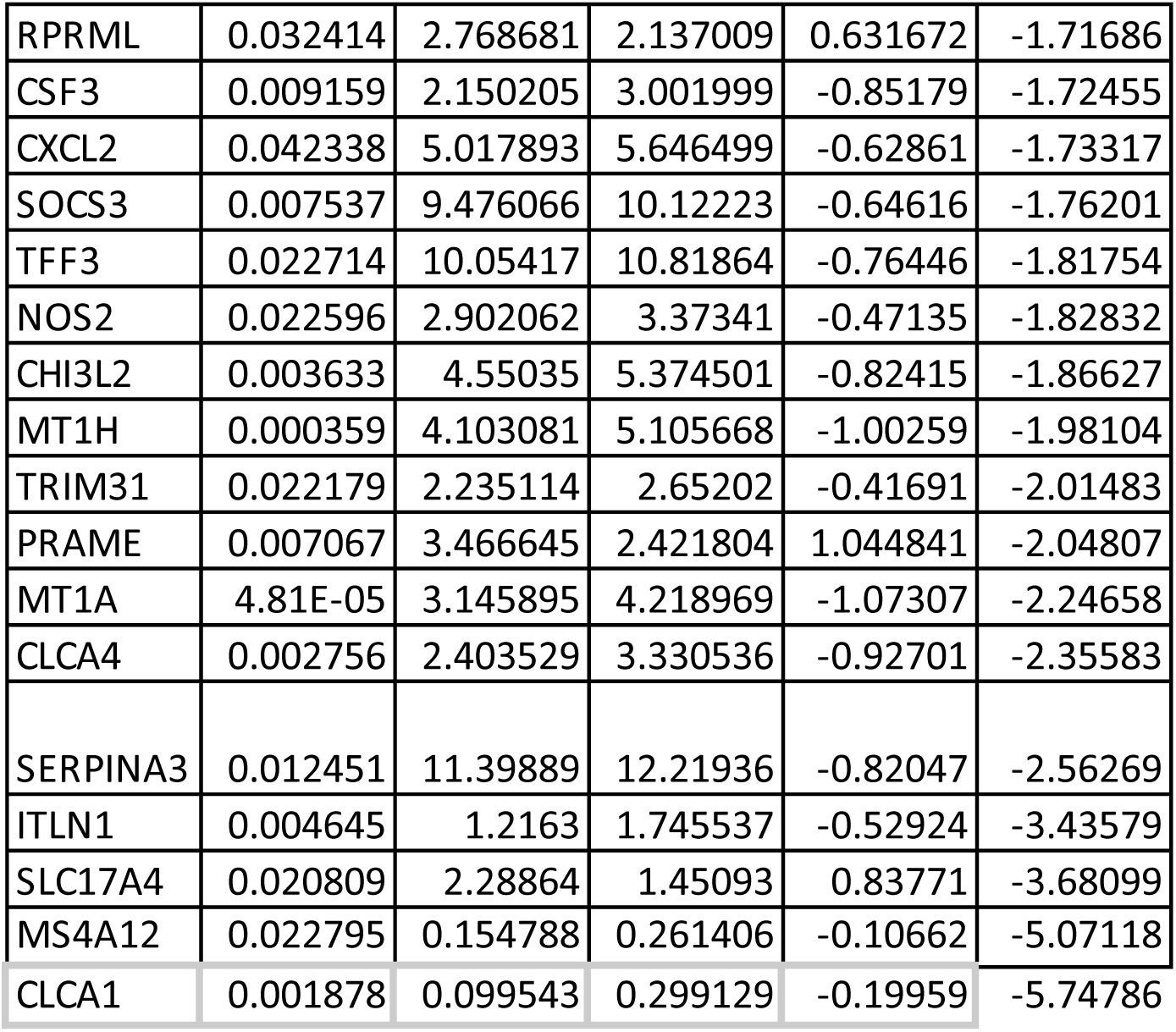
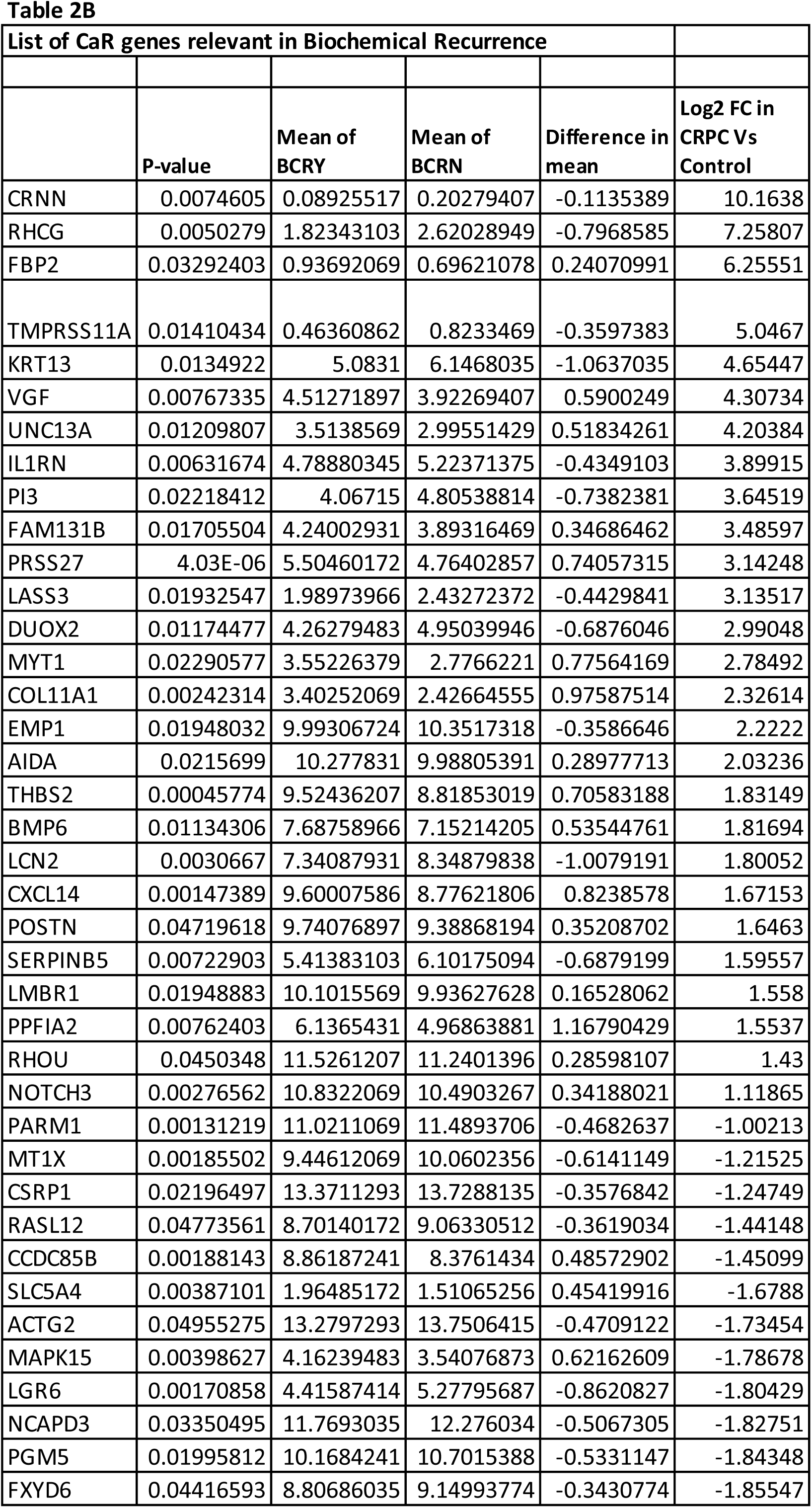
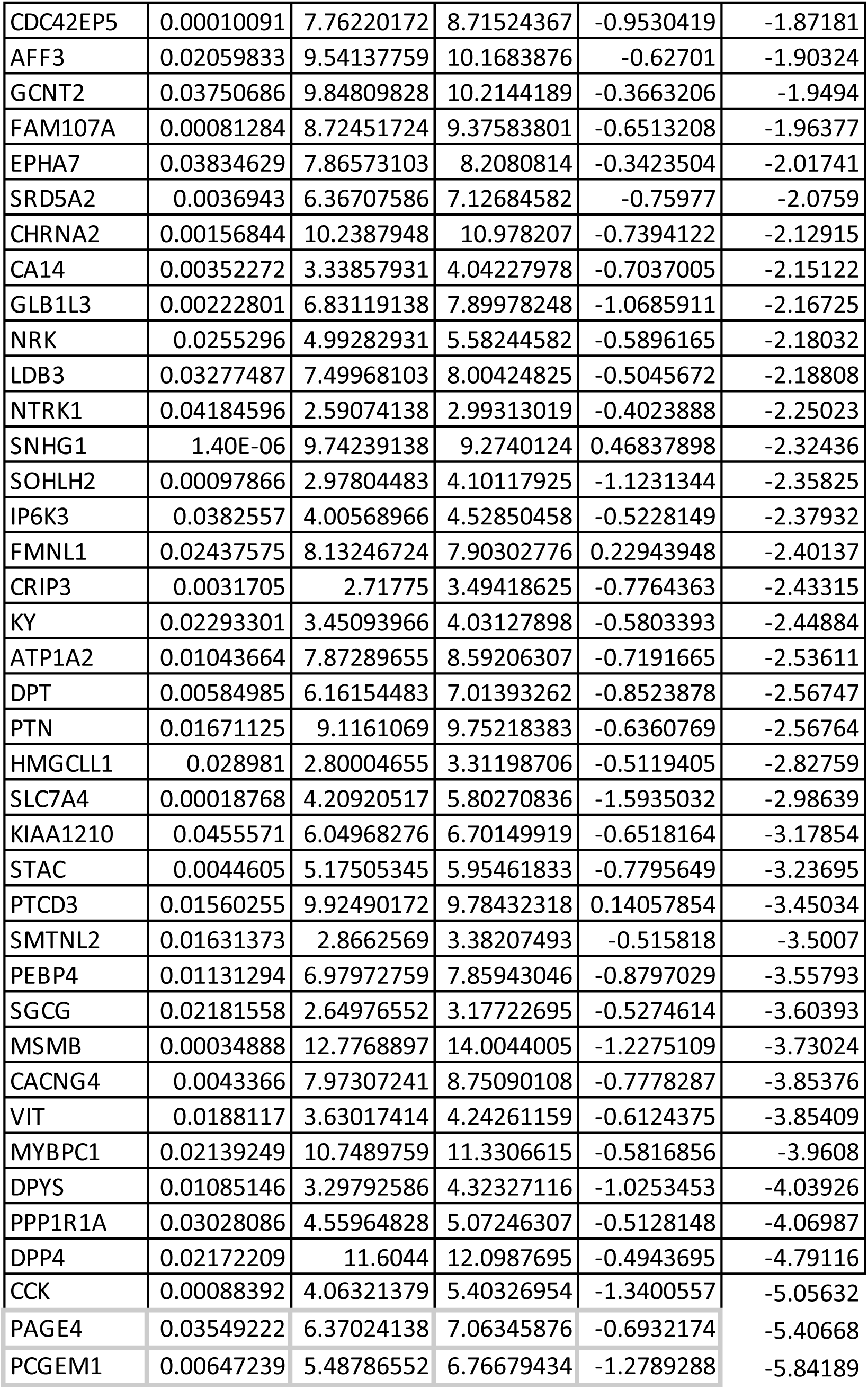
A. List of CaS genes involved in BCR (56) B. List of CaR genes involved in BCR (78) Note: The columns BCRY and BCRN give the average gene expression values in samples showing BCR and not showing BCR respectively. If the difference between these is positive, the gene is overexpressed in samples with BCR. These values are from TCGA data.

### Identification of genes associated with Gleason Score

Gleason grading is an established prognostic indicator of the aggressiveness of prostate cancer. Gleason score is assigned based on the histologic pattern of arrangement of cells in H&E-stained tissue sections of prostate carcinoma [20]. Correlation analysis of CaS genes with Gleason Score showed 160 CaS genes were significantly correlated with the Gleason Score with spearman rho value ranging from 0.37 to -0.38. Among them, 80 genes were positively correlated, and 80 genes were negatively correlated (Table 3A). Further, 191 CaR genes were significantly correlated with the Gleason Score with spearman rho value ranging from 0.38 to -0.46. Among them, 53 genes were positively correlated, and 138 genes were negatively correlated (Table 3B).

**Table 3:**
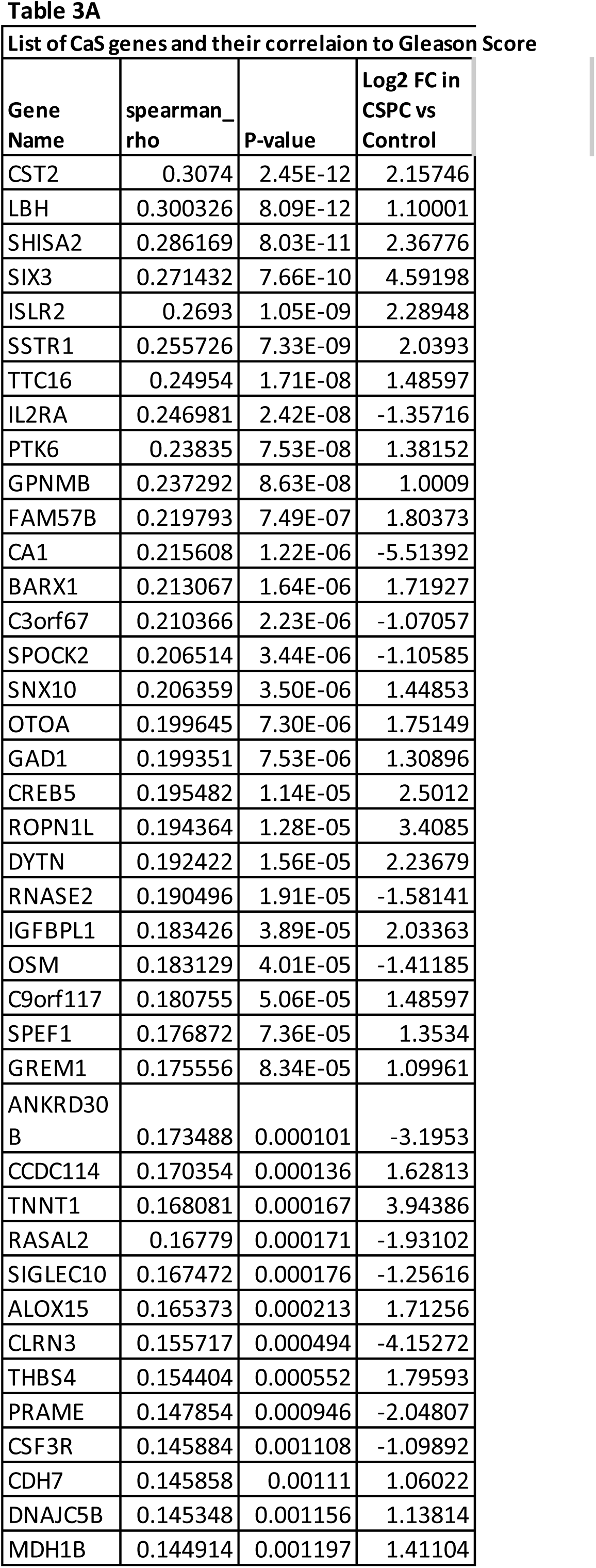
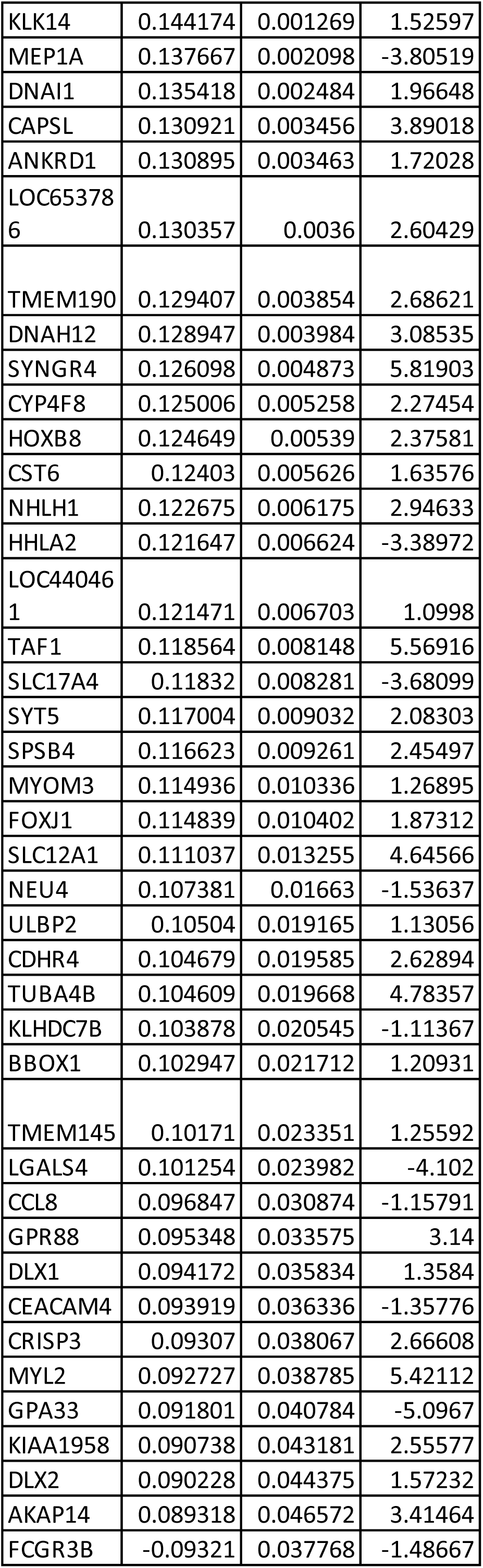
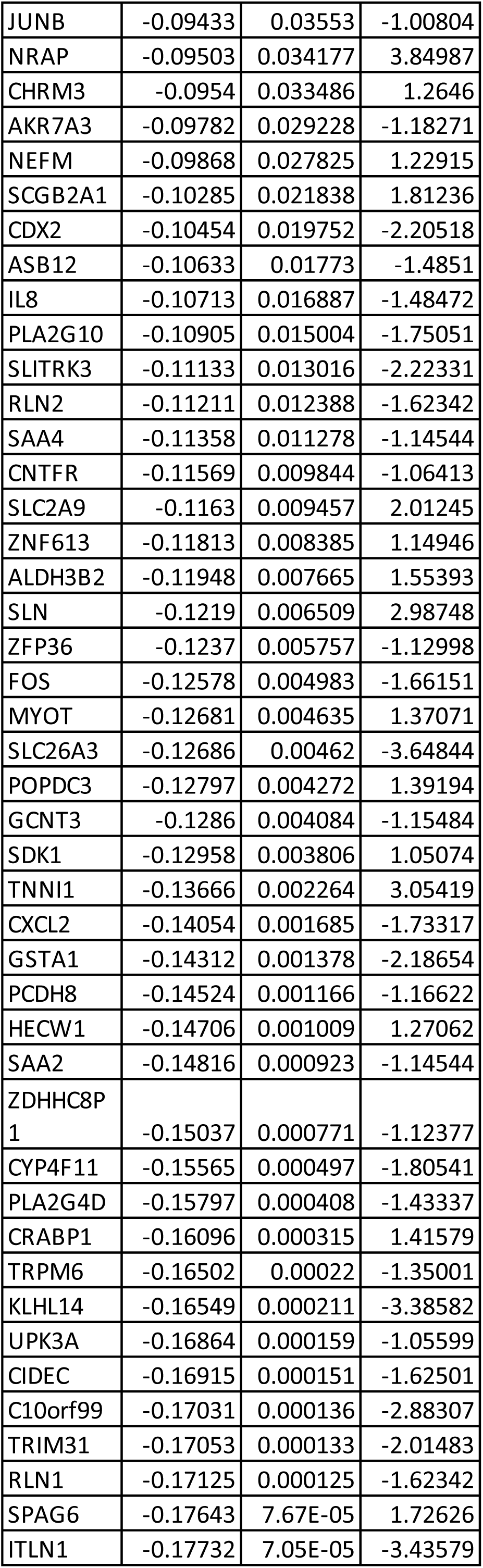
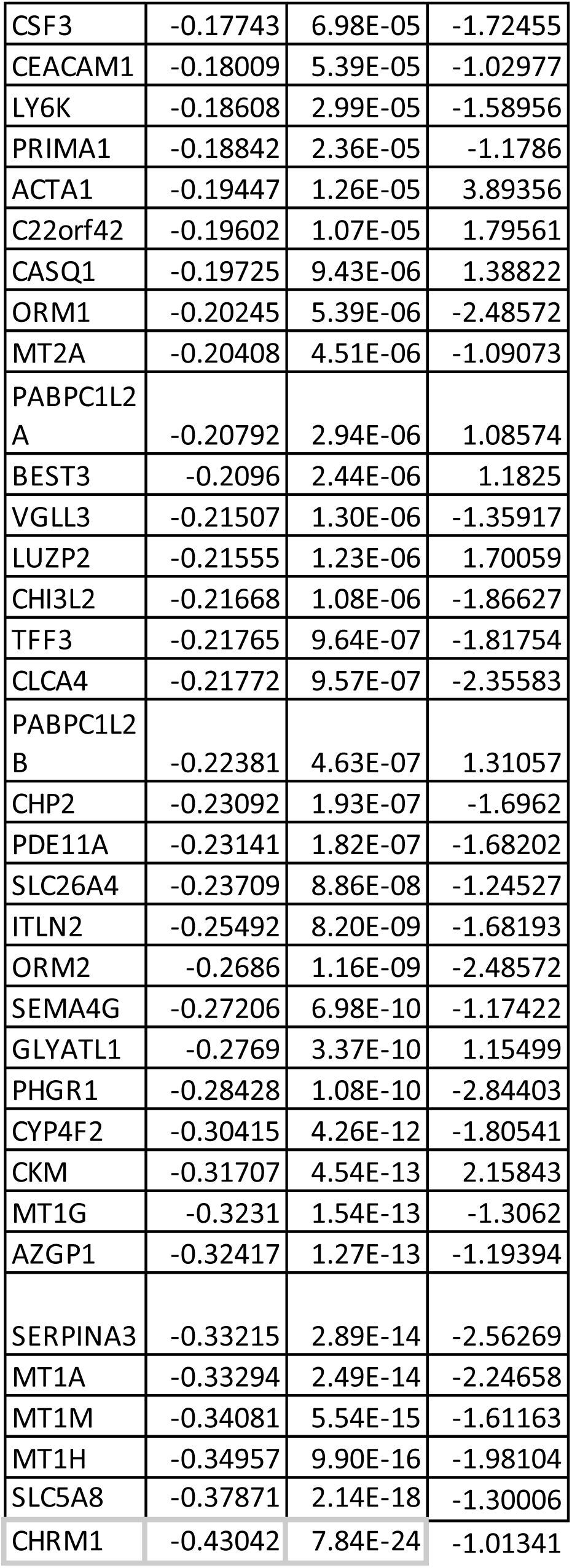
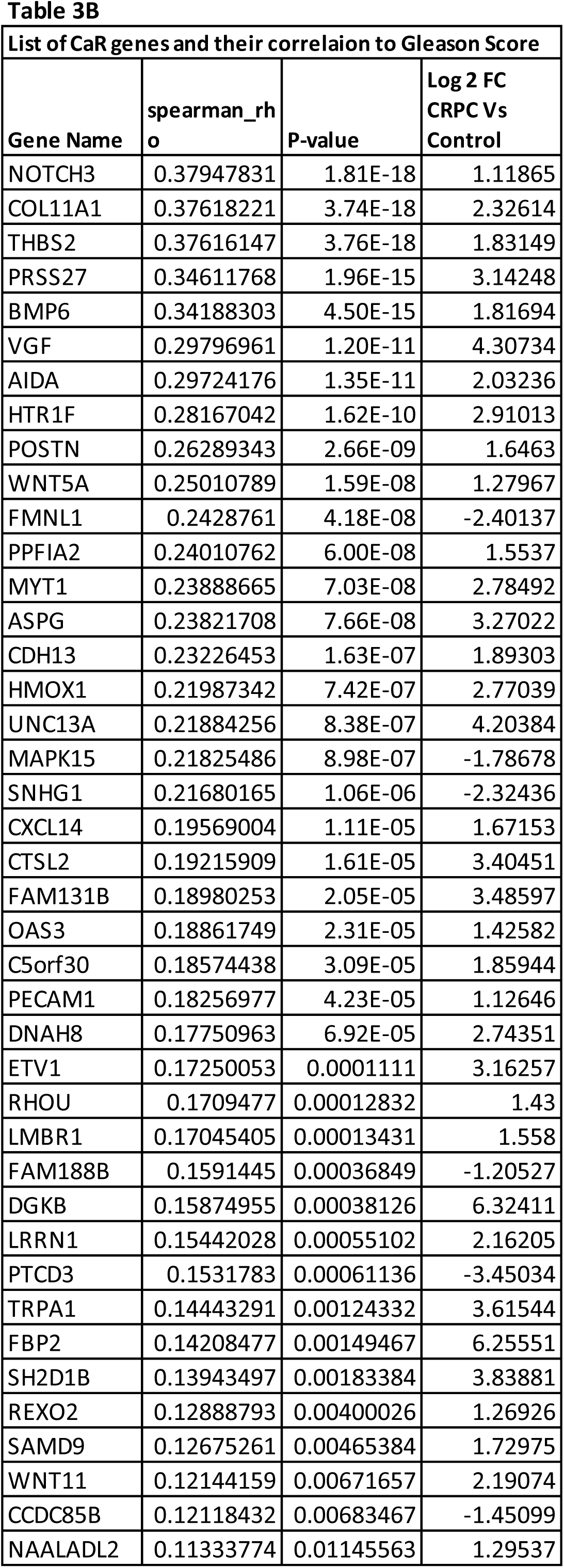
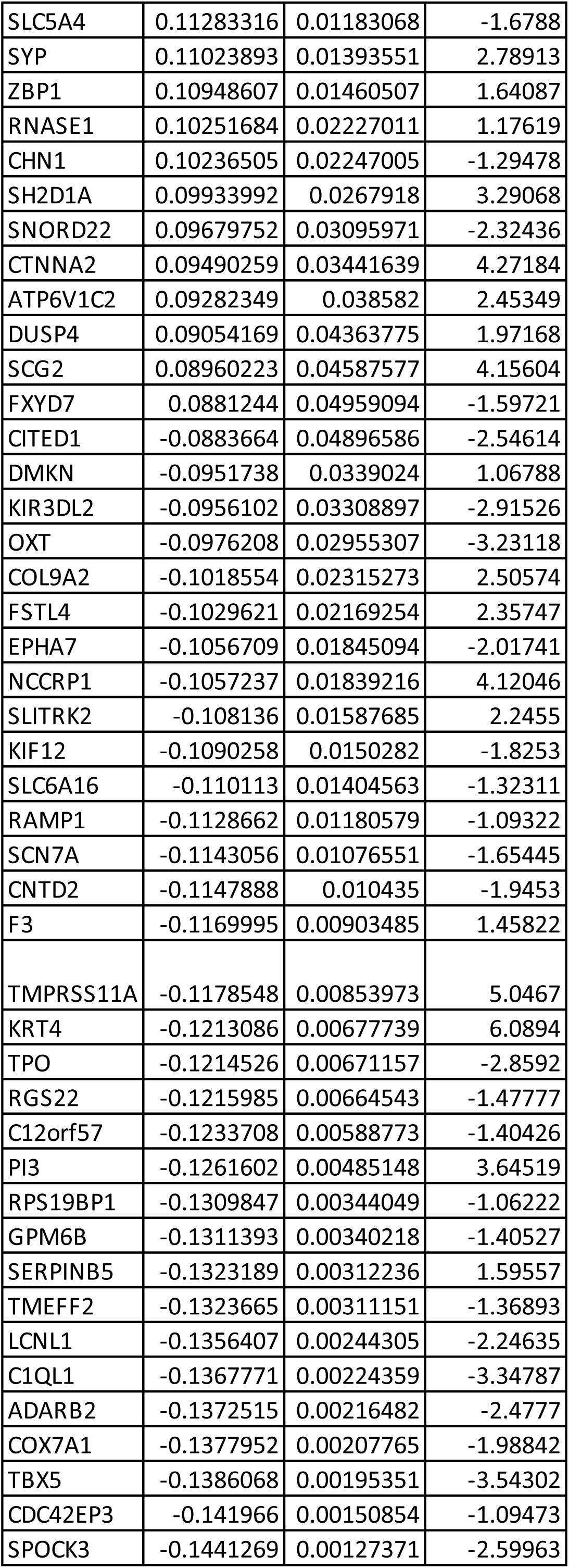
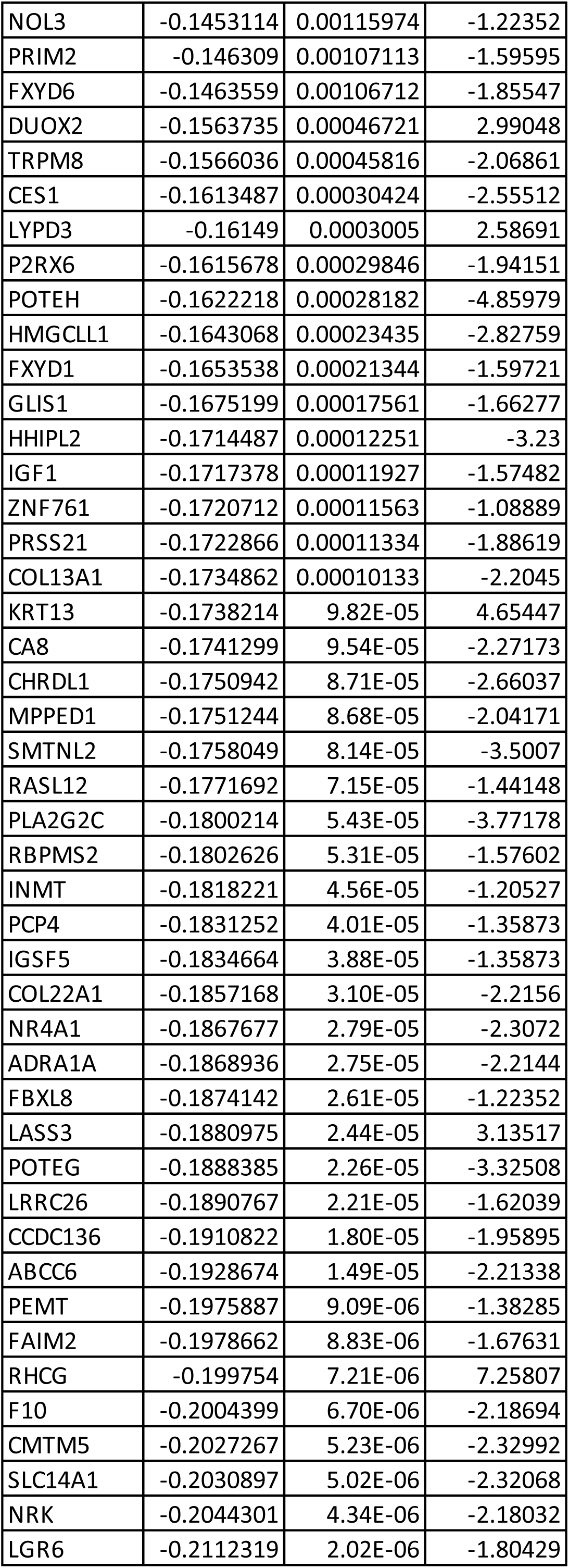
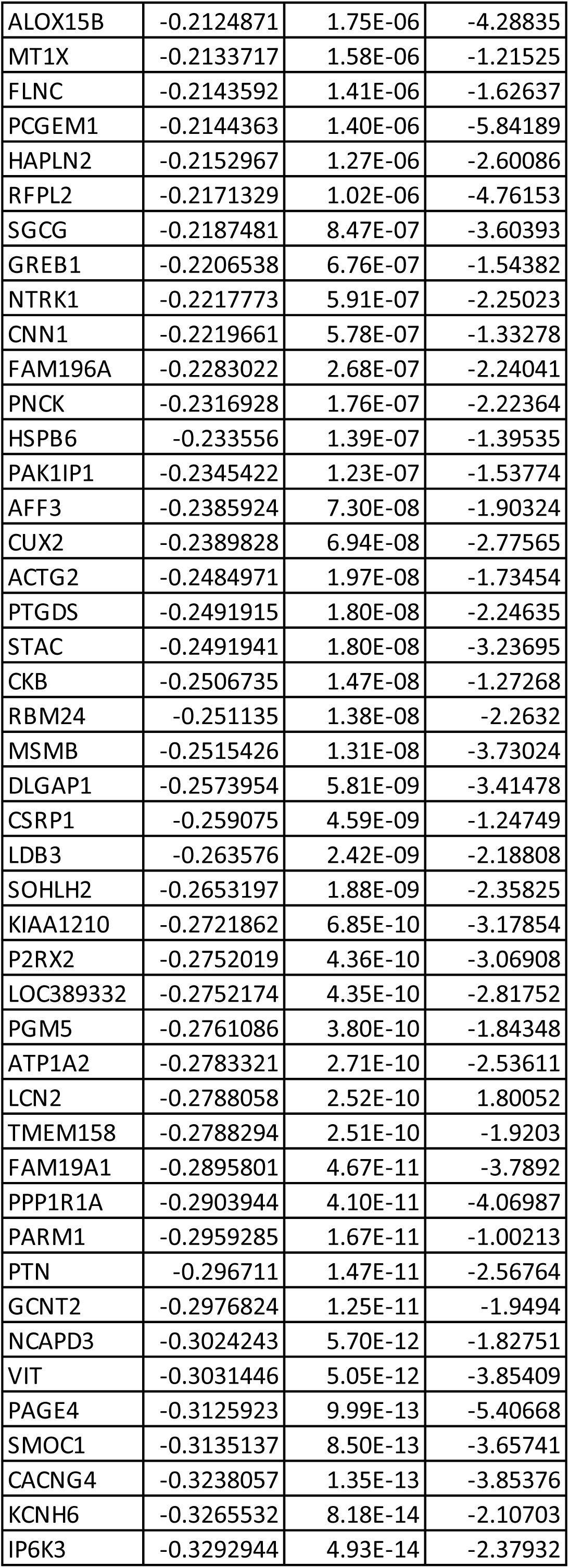
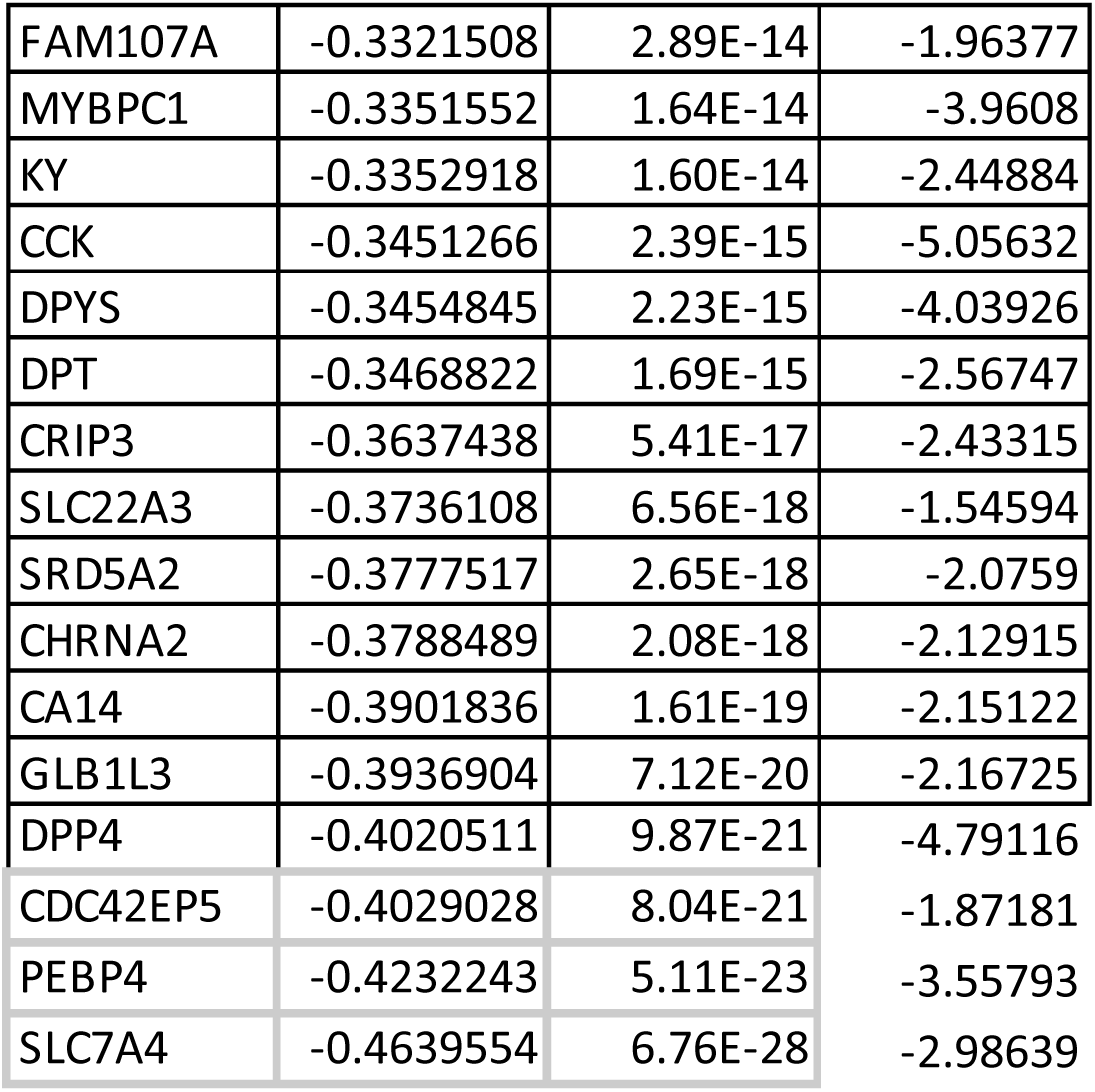
A. List of CaS genes and their correlation with Gleason Score (160) B. List of CaR genes and their correlation with Gleason Score (191)

Based on this data, it appears that the Gleason score may or may not have a major implication in development of CRPC.

### Identification of genes associated with tumor recurrence

To identify the CaR genes associated with the tumor recurrence that occurs after treatment, mRNA expression values of CaR genes in the samples with tumor recurrence and samples without tumor recurrence were analyzed. 99 CaS genes were differentially expressed in between the two groups (p < 0.05). 53 CaS genes showed more expression in samples with tumor recurrence, and 46 genes showed less expression in samples with tumor recurrence compared to samples without tumor recurrence (Table 4A). Also, 125 CaR genes were differentially expressed between the two groups (p < 0.05). 37 CaR genes showed more expression in samples with tumor recurrence, and 88 genes showed less expression in samples with tumor recurrence compared to samples without tumor recurrence (Table 4B).

**Table 4:**
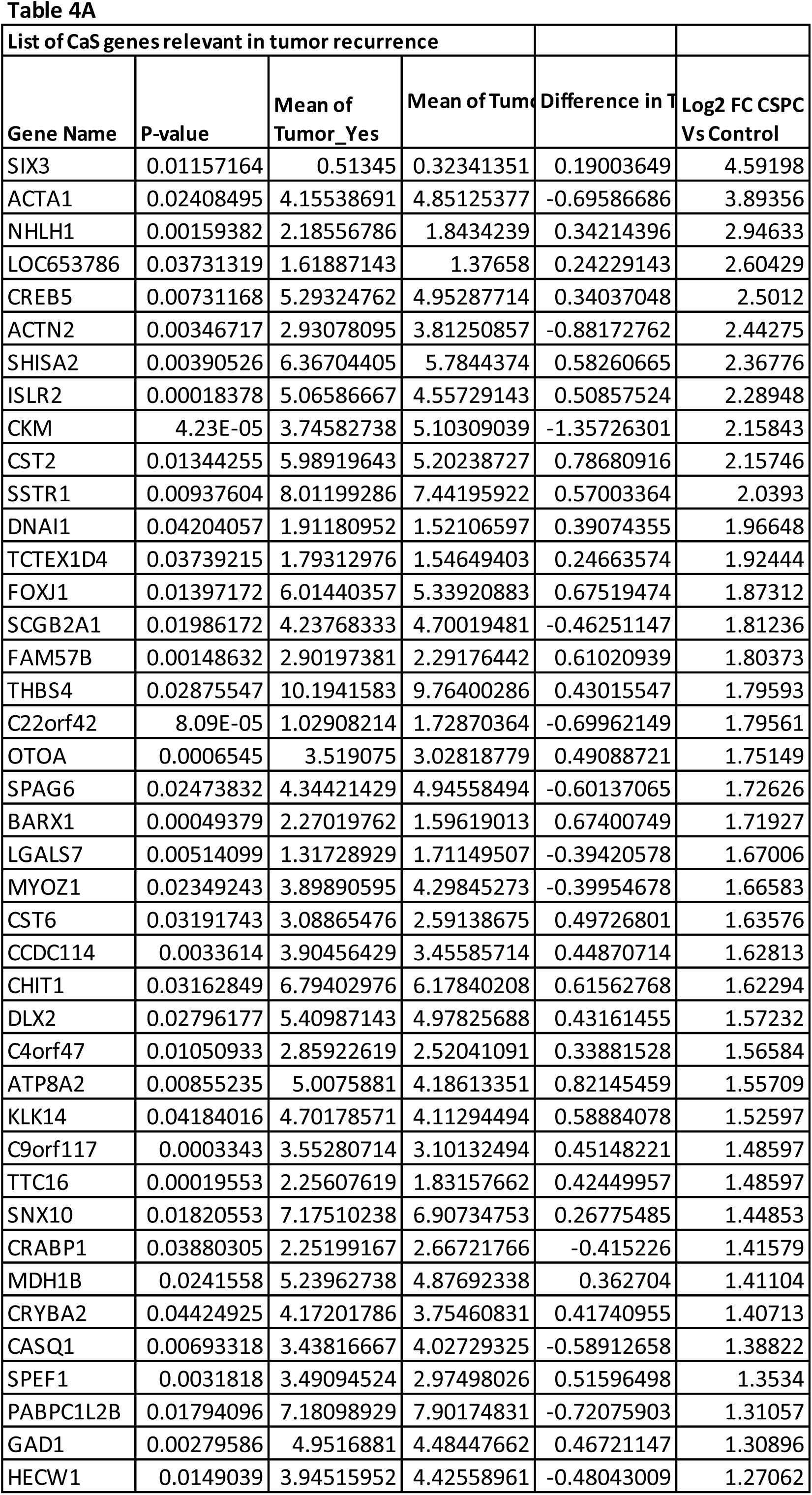
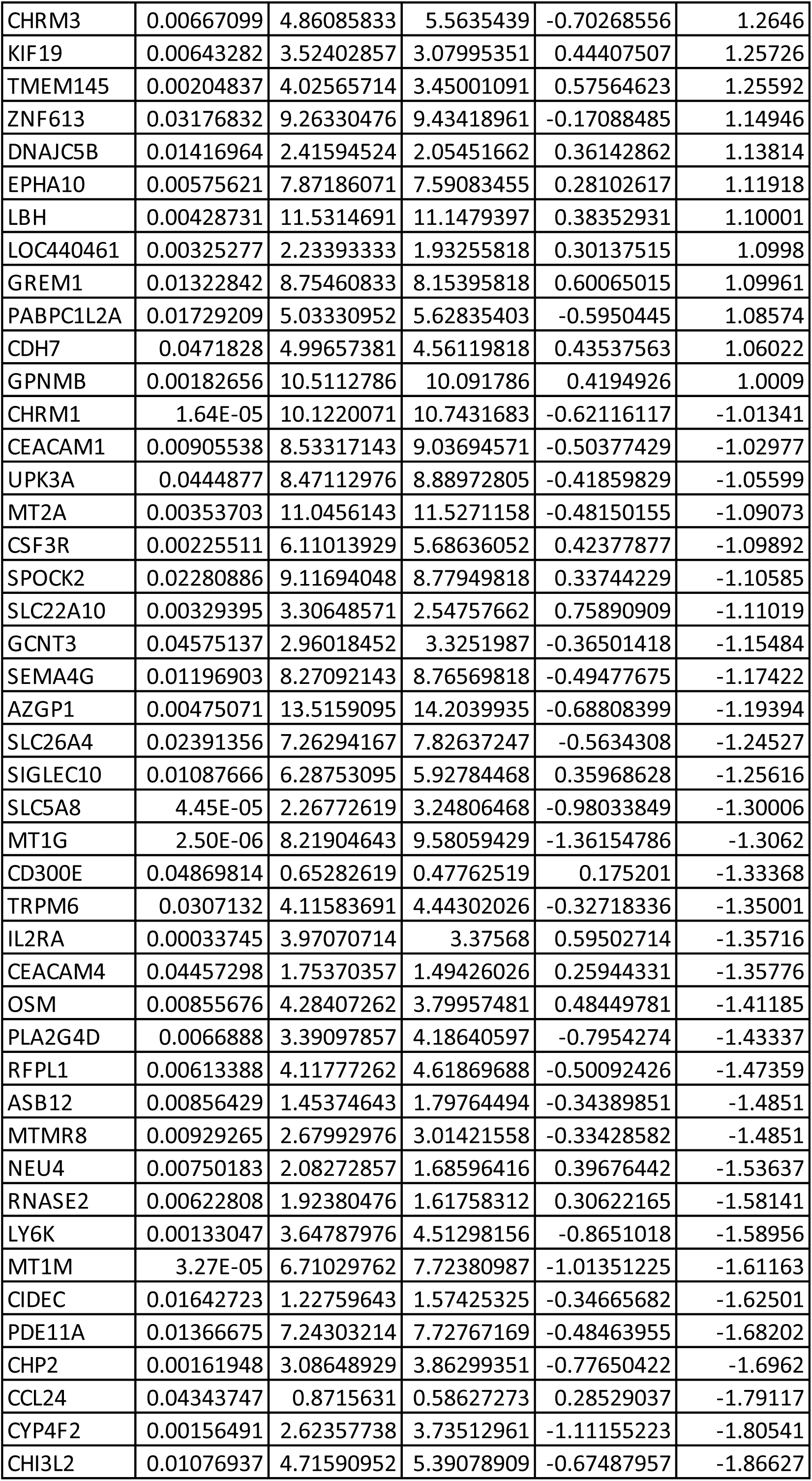
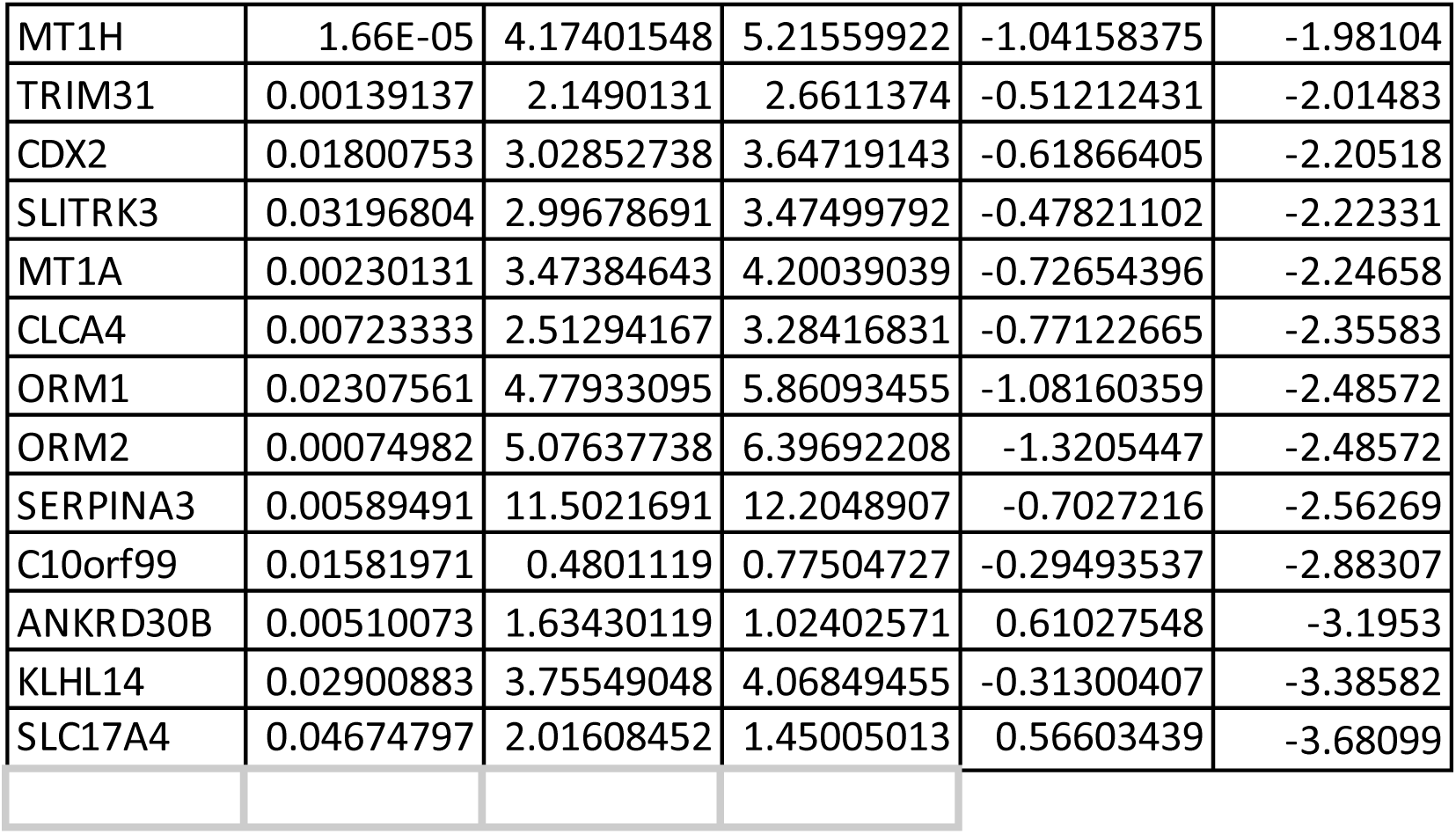
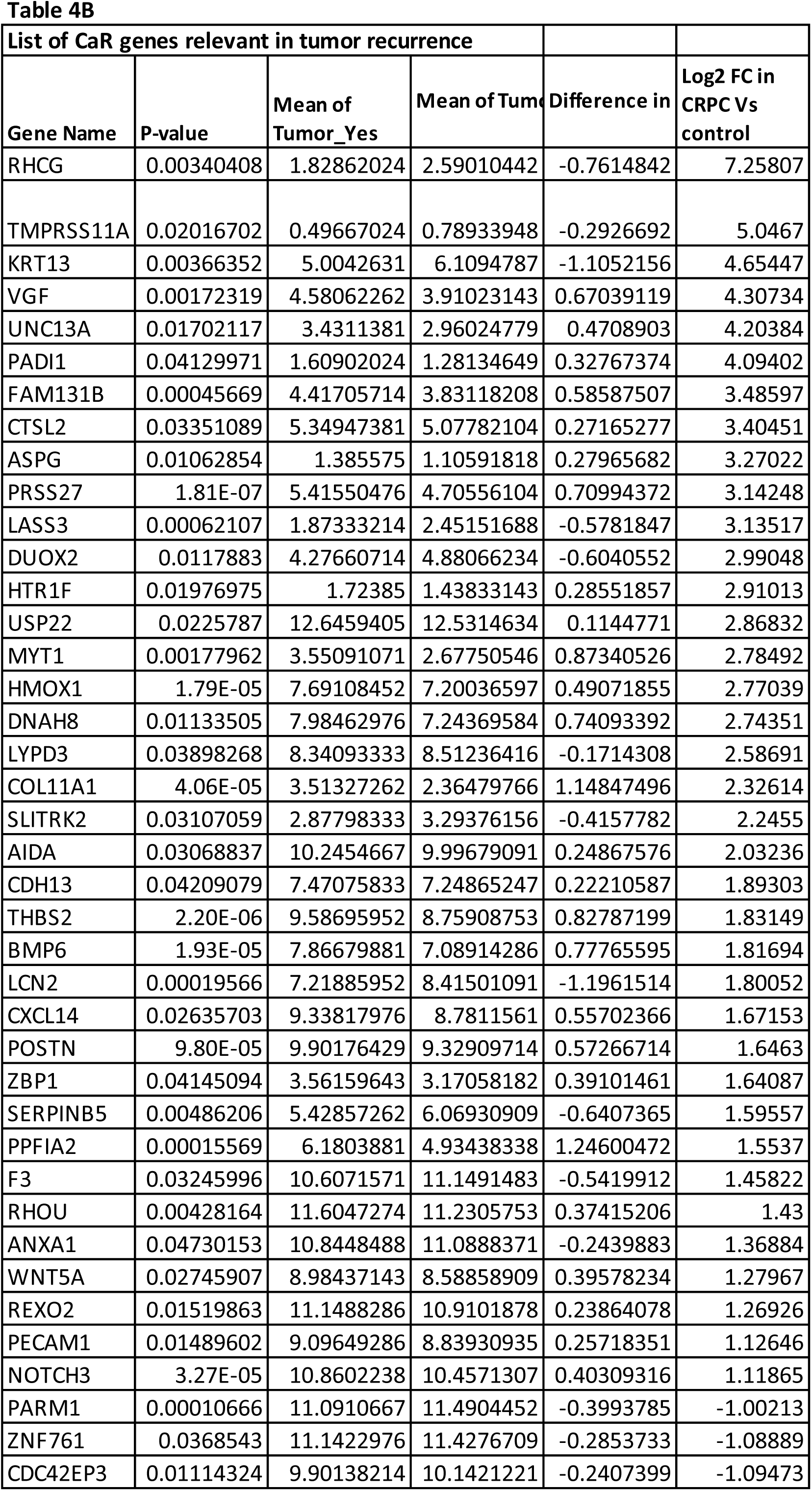
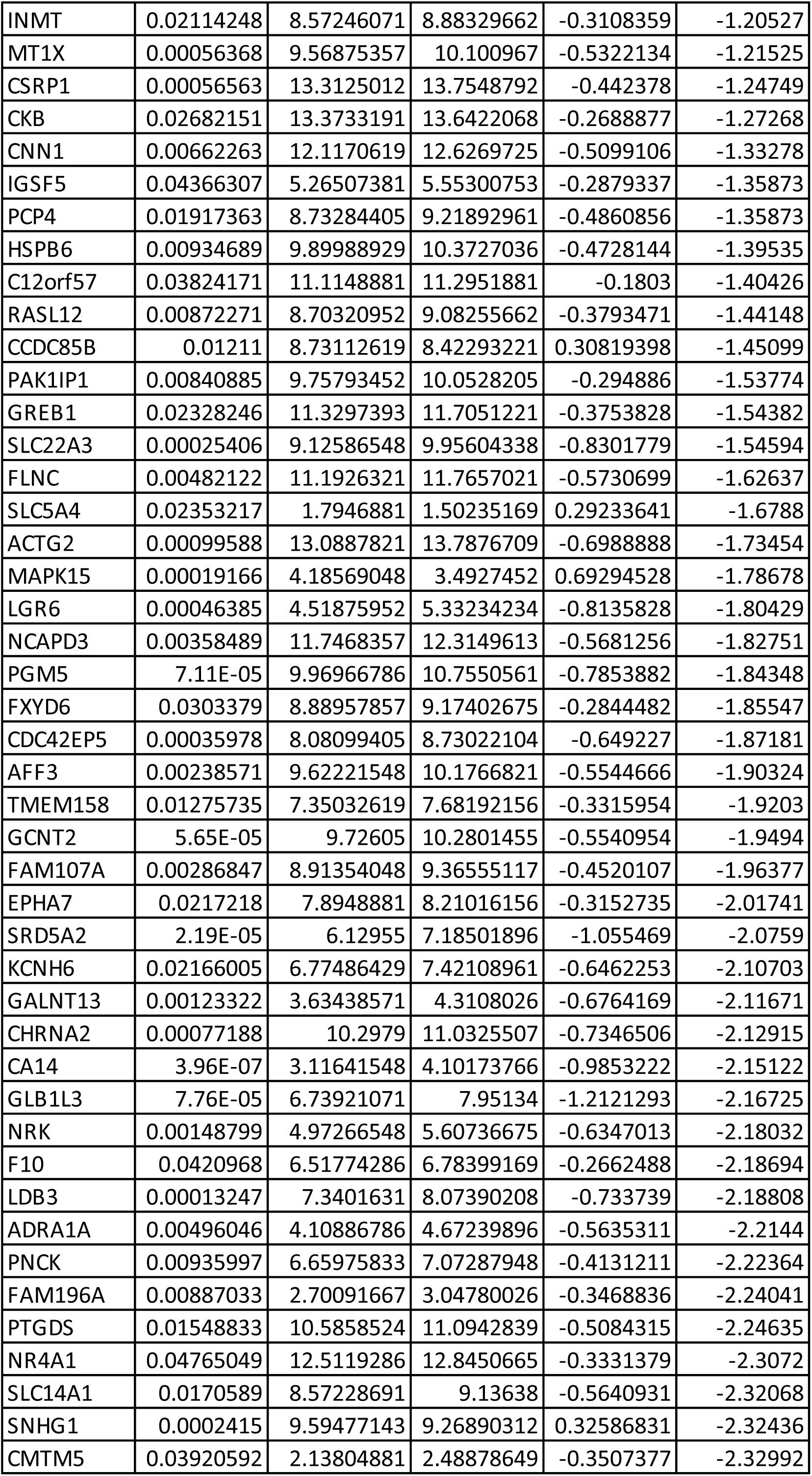
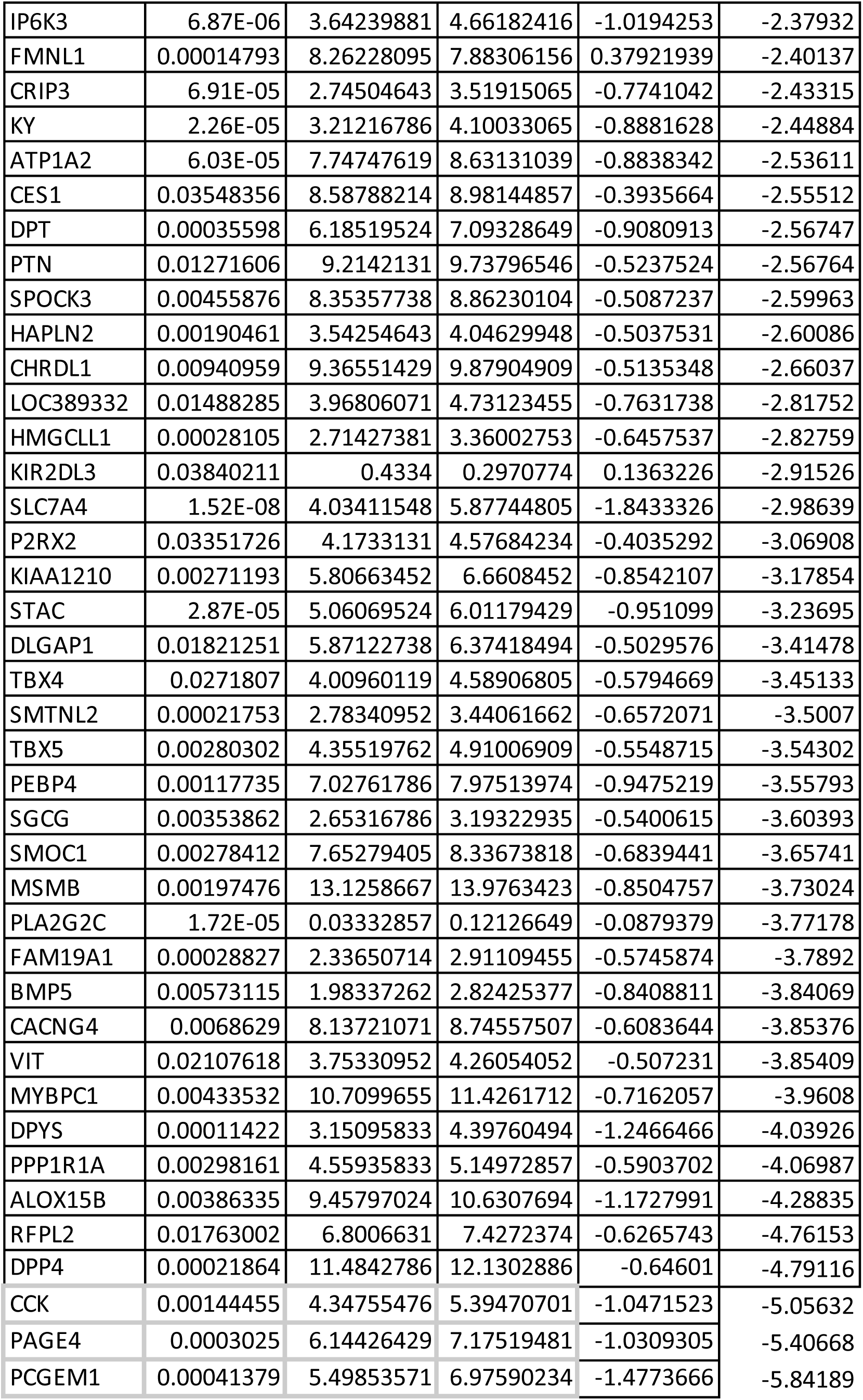
A. List of CaS genes which are differentially expressed between recurrent and non-recurrent groups (99) B. List of CaR genes which are differentially expressed between recurrent and non-recurrent groups (125) Note: The columns Tumor Y and Tumor N give the average gene expression values in samples showing recurrence and not showing recurrence respectively. If the difference between these is positive, the gene is overexpressed in samples with recurrence. These values are from TCGA data.

### Analysis of prognostic significance of the CaR genes and CaS genes

To evaluate the prognostic value of the CaS and CaR genes, Kaplan Meier survival analysis of CaR genes was done using TCGA survival data (Figure 2, Supp fig 3). Out of 292 CaR genes, 9 CaR genes were significantly associated with overall survival (with a range of hazard ratios (HR) 0.12–10.17 and p-values 0.0071 to 0.049), 96 genes were associated with progression-free interval, 69 genes were associated with disease-free interval (Table 5B). Further, among 373 CaS genes, 13 genes were significantly associated with overall survival, 69 genes were associated with progression-free interval, 47 genes were associated with disease-free interval (Table 5A). Representative survival plots for select genes in each of these categories is depicted in Figure 2.

**Figure 2:**
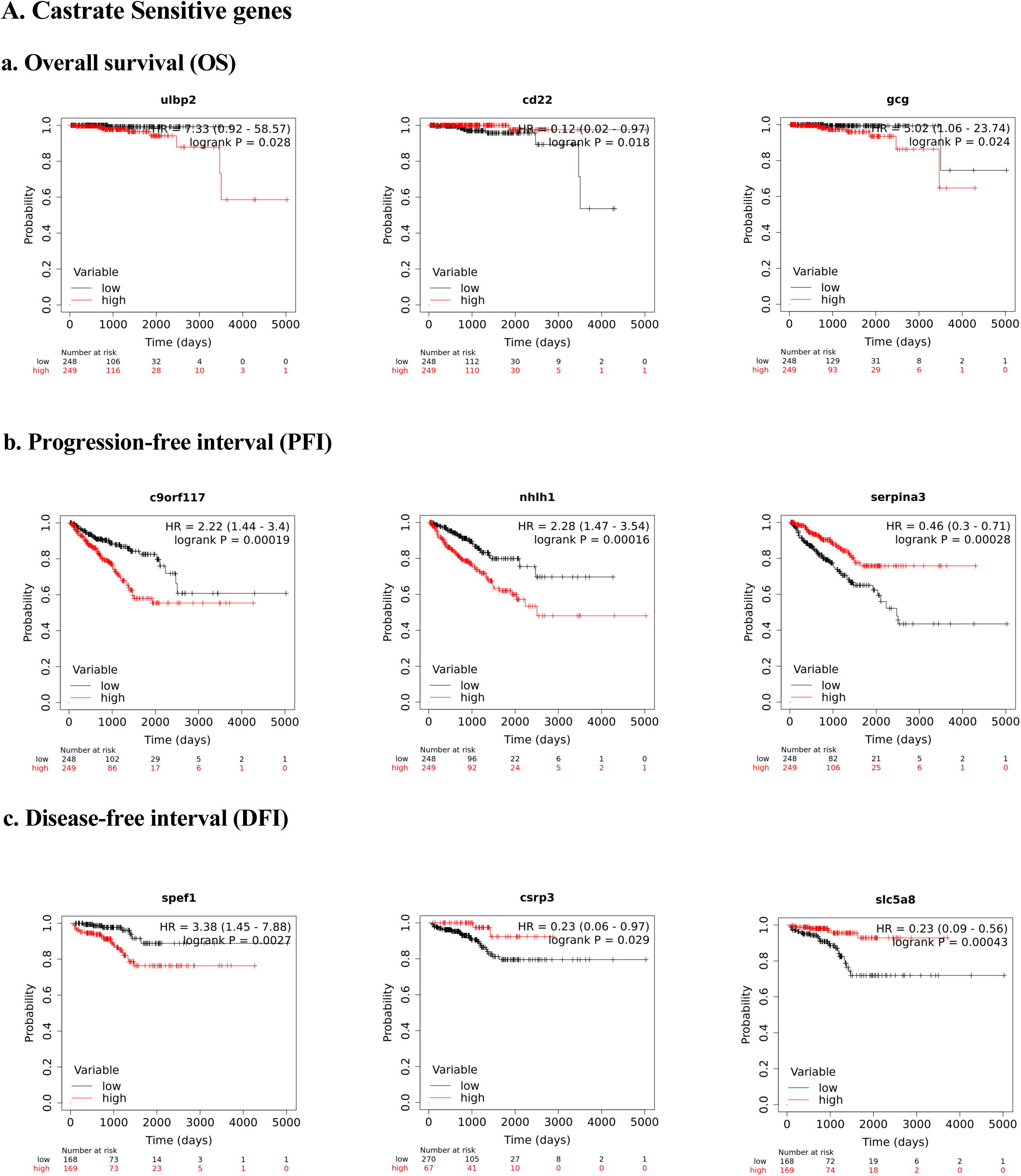
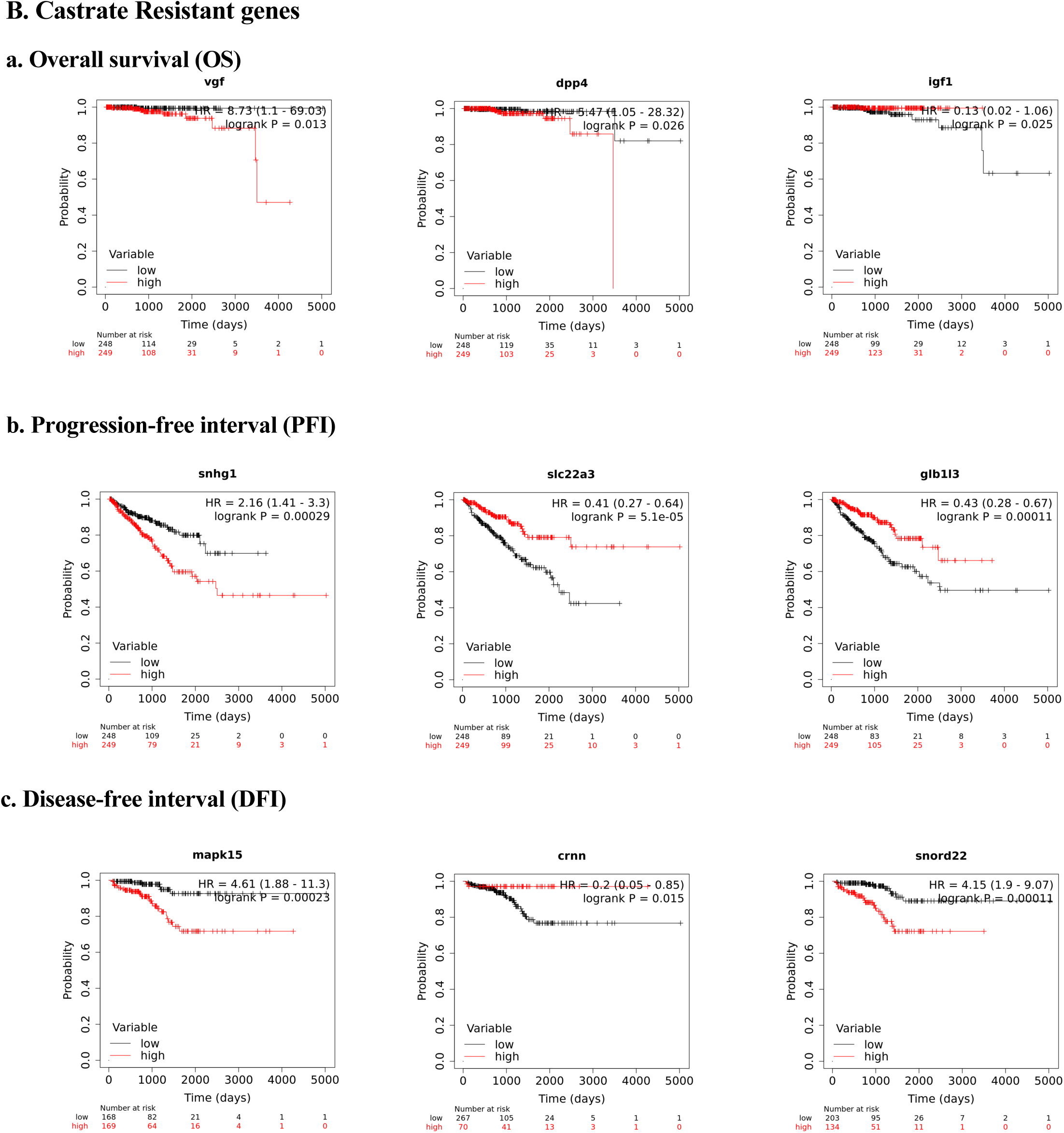
A. Shows the K-M analysis of selected CaS genes and their impact on a. Overall survival b. Progression free survival c. Disease free progression B. Shows the K-M analysis of selected CaR genes and their impact on a. Overall survival b. Progression free survival c. Disease free progression (P.S: The analysis for other genes can be made available upon request)

**Table 5:**
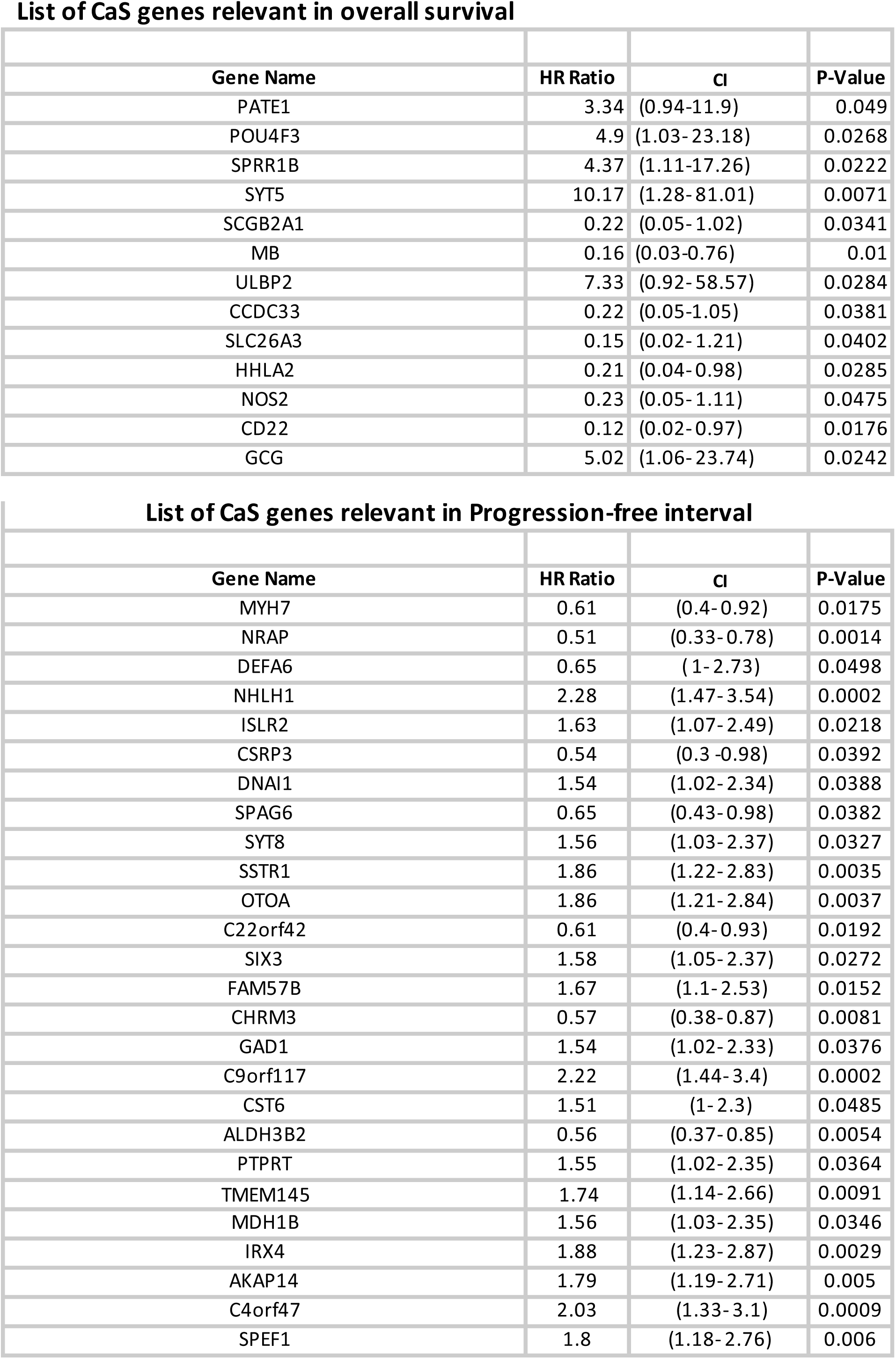
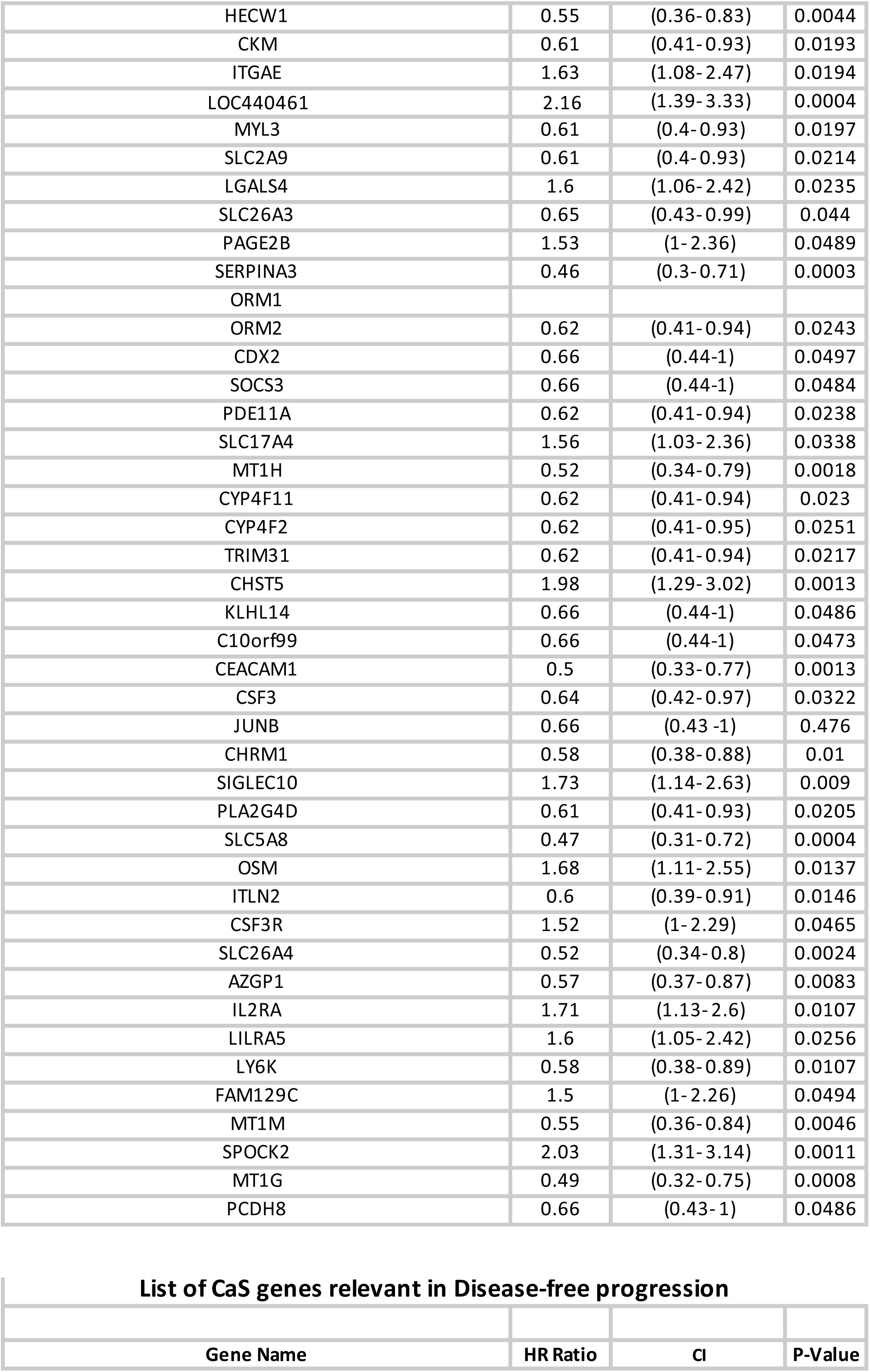
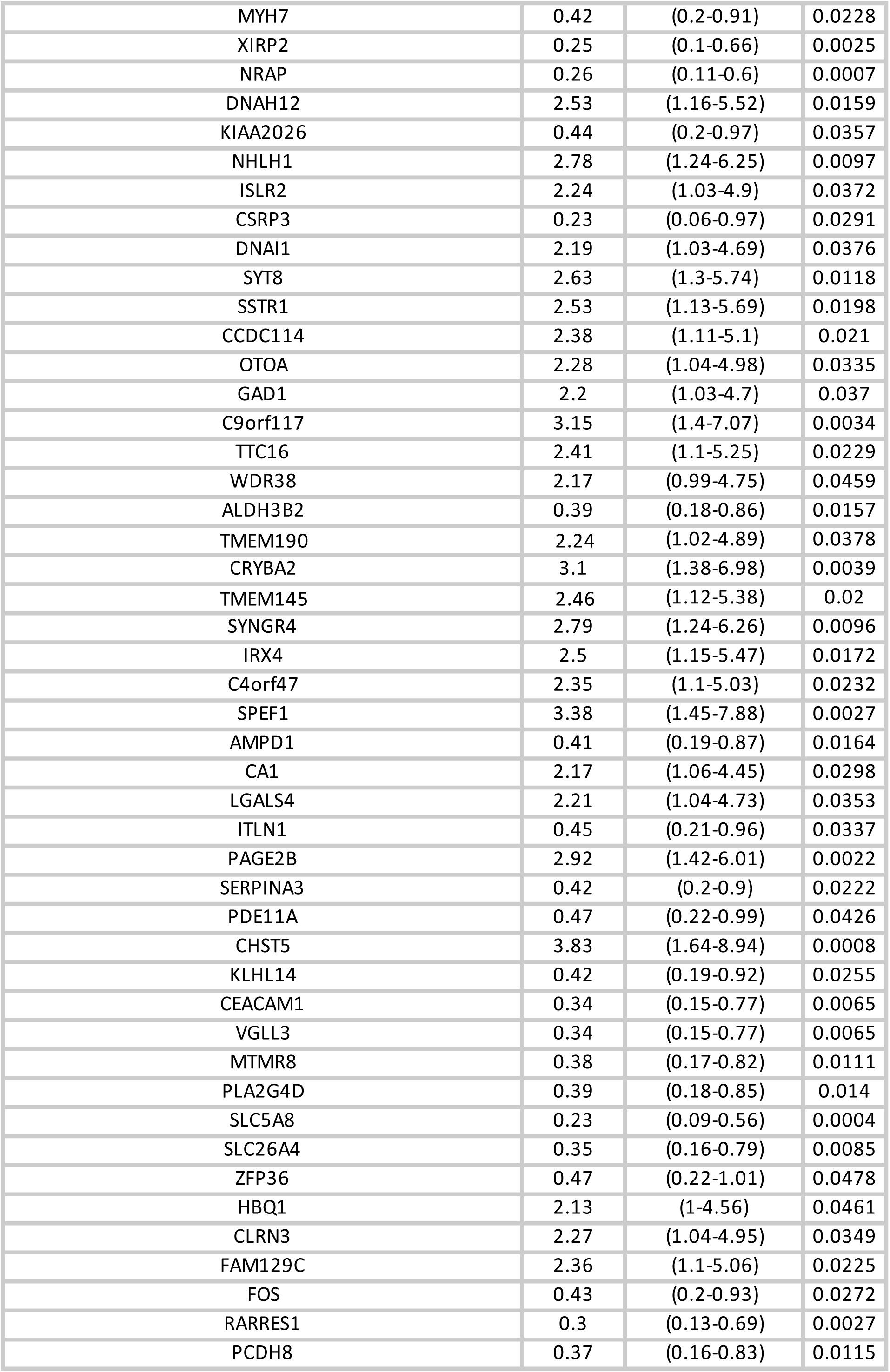
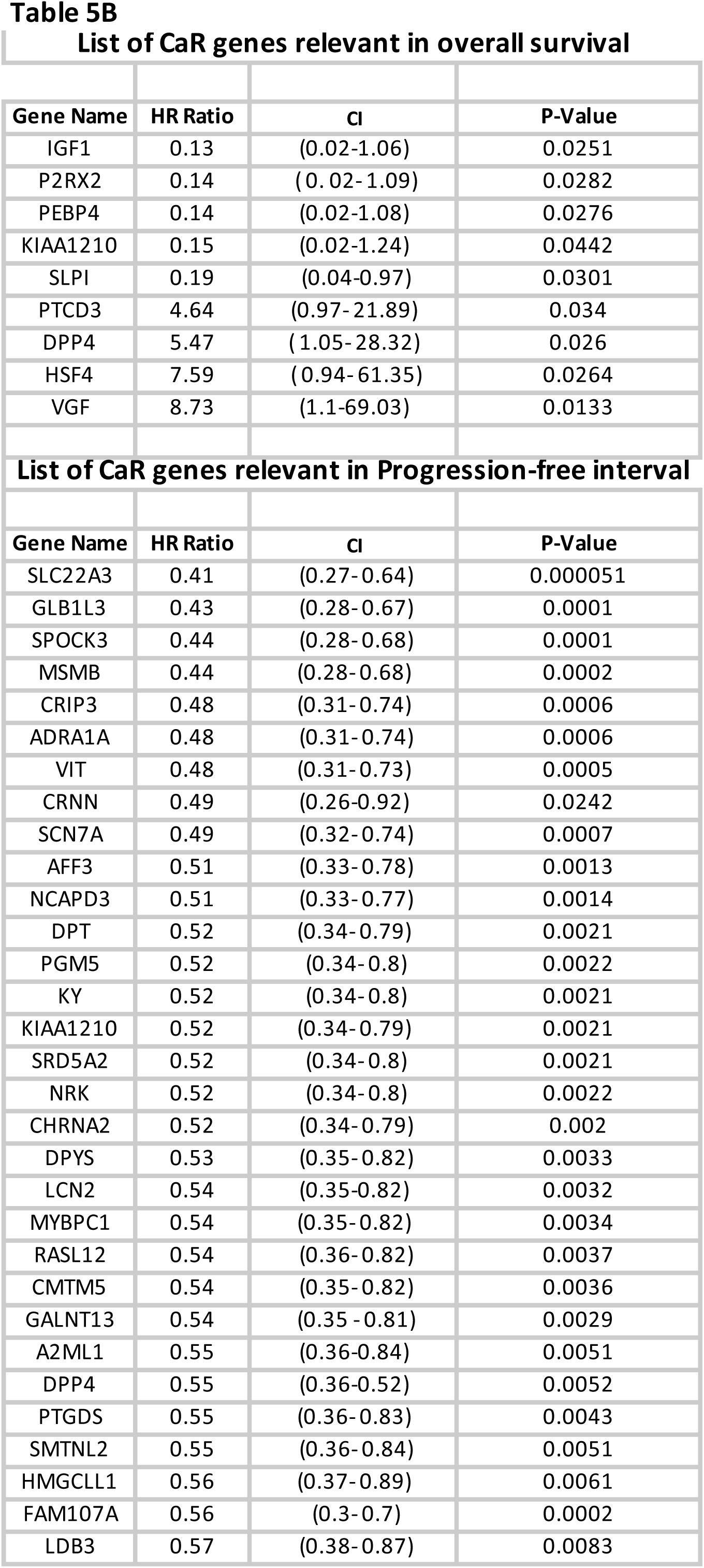
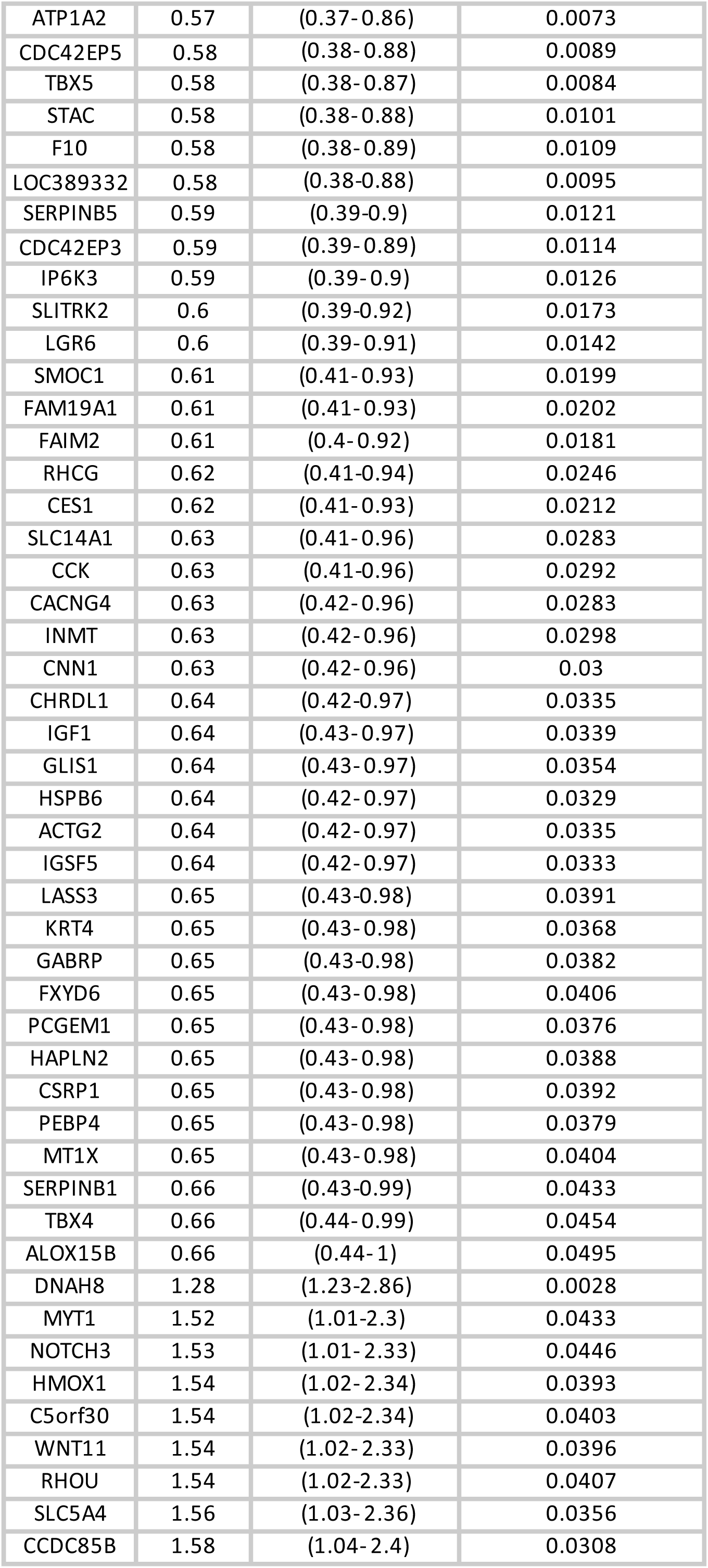
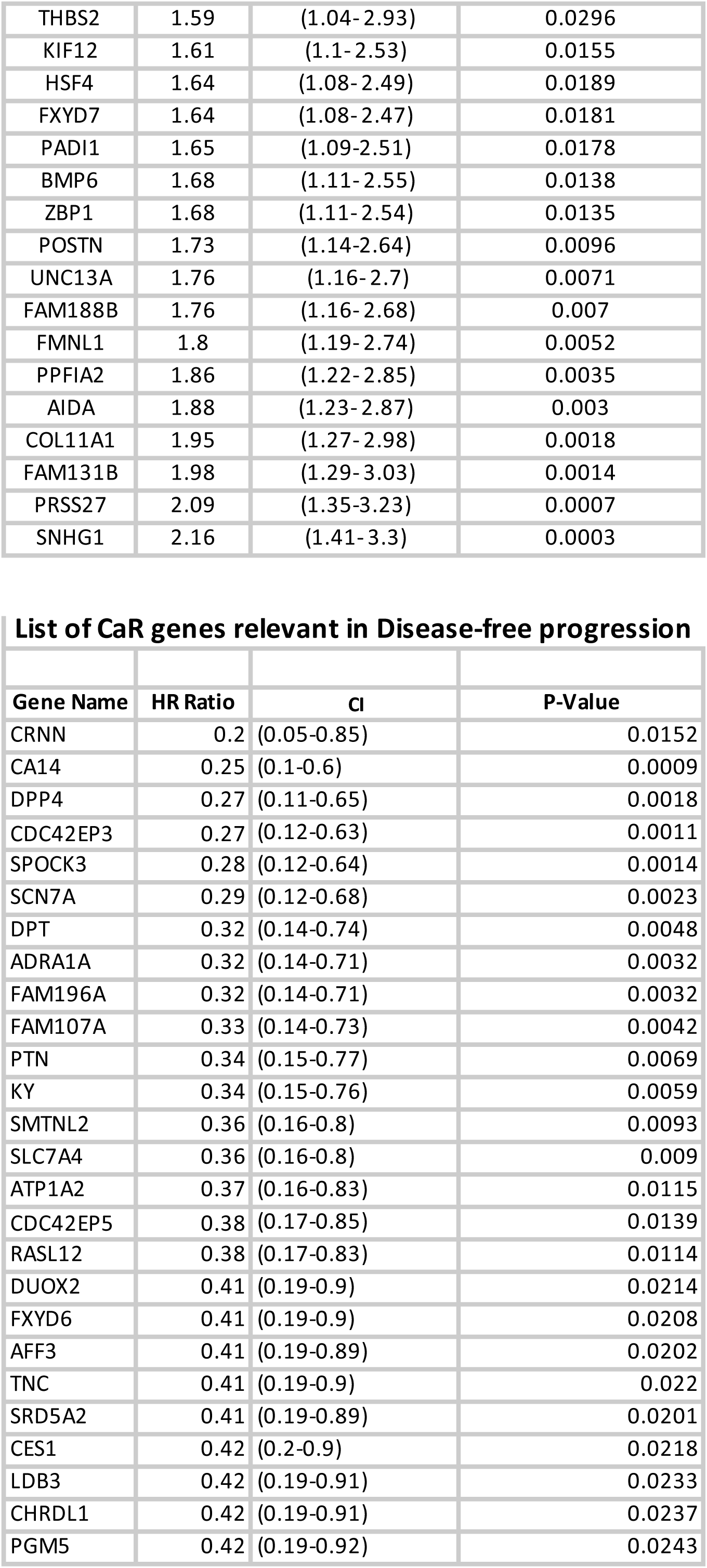
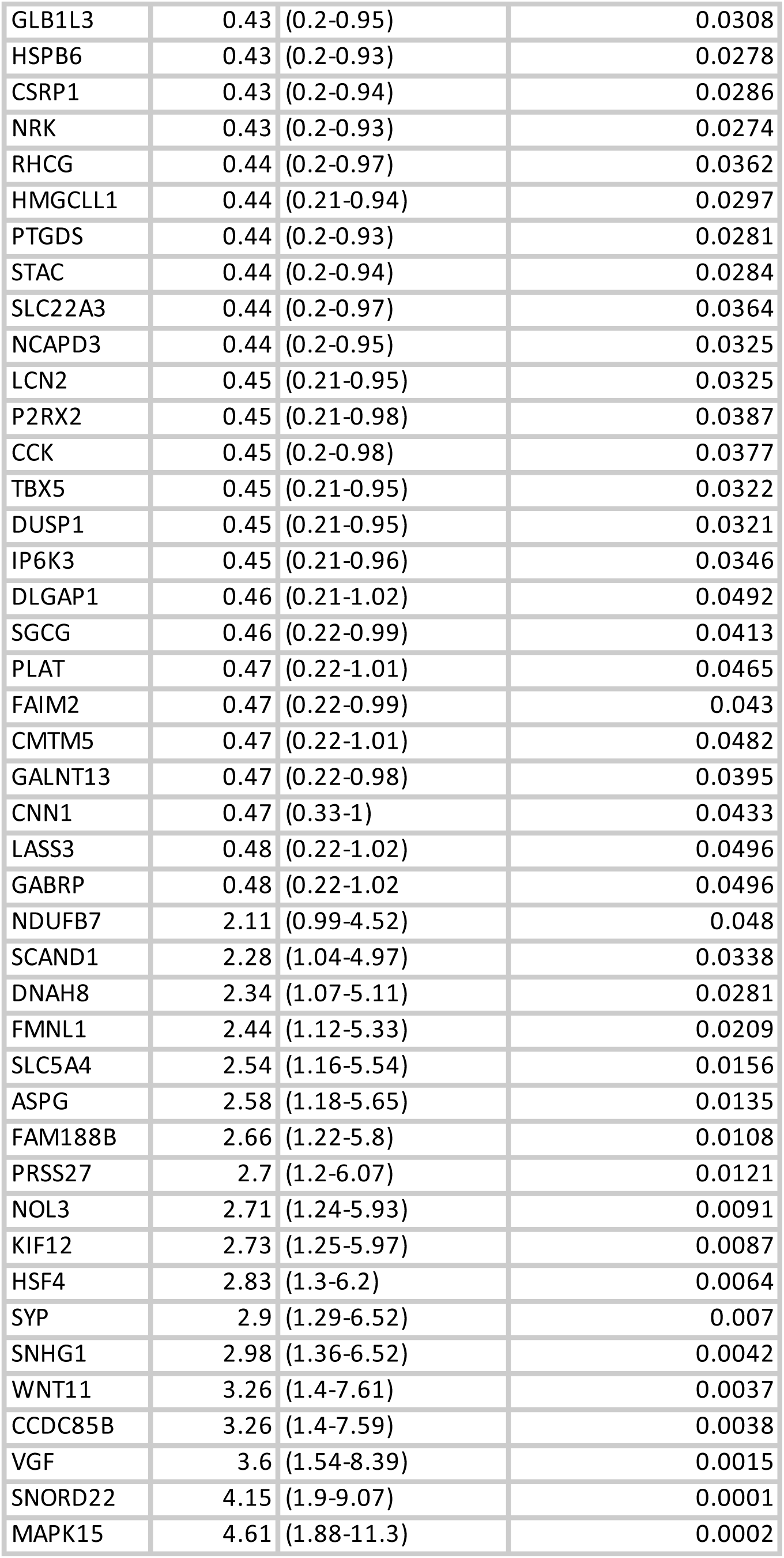
A. List of CaS genes which are associated with i. Overall survival, ii. Progression free survival, iii. Disease free progression B. List of CaR genes which are associated with i. Overall survival, ii. Progression free survival, iii. Disease free progression

Although the number of CaR genes associated with overall survival is not very significantly different from the number of CaS genes, there seems to be a difference in the genes which may impact progression of the tumor, namely progression free interval and disease free interval. However this requires validation on a larger sample size as well as mechanistically.

## Discussion

Given that prostate gland is highly dependent on androgen signaling for development as well as normal functioning, androgen deprivation therapy is a widely used treatment option particularly in metastatic prostate cancer. However, despite being effective, sooner or later, majority of the patients develop resistance to ADT, following which the prognosis is poor.

Many mechanisms that may be responsible for the development of this resistance have been proposed. Several groups have identified genetic and epigenetic alterations in genes such as Rb1 and beta catenin [9] and also Wnt pathway genes such as CTNNB1, APC or RNF43 which can lead to aberrant activation of the pathway [10]. Many of these markers would serve as excellent markers for identifying androgen resistance and also for prognosis. However, considering the fact that most of these mutations would eventually lead to aberrant androgen signaling or cross talk with other pathways, they would also lead to changes in gene expression patterns. Identifying changes in the gene expression, both qualitative and quantitative would not only serve as markers of prognosis, but also may serve as targets for therapeutic intervention to overcome the problem of resistance.

In this study, we have profiled gene expression patterns of 23 castrate sensitive and 4 castrate resistant prostate cancers and 31 BPH samples. We have identified genes which are differentially expressed between BPH and CaS, BPH and CaR as well as CaS and CaR groups. Based on these comparisons, we have classified the genes as CaS genes and CaR genes. We do acknowledge that this classification may not be very accurate given the small number of samples in the CaR group. However, they would be of value as a proof of concept. In order to check if these genes have any relevance in the prognosis, we have compared our profiles with data from TCGA [15]. TCGA has gene expression data from 498 prostate cancer patients and 52 controls. However, information on whether these are CaS or CaR is not available. Hence these two sets of data can complement each other. For all the analyses done with respect to relevance of genes, we have used only those genes which were present both in our data and TCGA.

What we have tried to correlate is whether the CaR or CaS gene expression signature correlates to any prognostic parameters. The parameters we have taken into consideration are biochemical recurrence, Gleason score, tumor recurrence and survival. In the case of biochemical recurrence and tumor recurrence, we have seen that more of the CaR genes seem to correlate with recurrence than CaS genes. It could imply that the CaR signature are likely to be predictive of recurrence than CaS signature. However, to make this correlation stronger, more validation would be required. When we consider the Gleason score, which is an indicator of aggressiveness of the tumor, there seems to be no correlation with the CaR or CaS genes.

The other prognosis indicator we have considered is Survival. In this, we have taken into consideration overall survival, progression free survival and disease free progression. The CaR signature doesn’t seem to impact overall survival. However, with the given data, it appears that the CaR signature seems to have an impact on the progression of the disease and also the quality of life for the patient. When we compare the gene expression signature which is common to all the parameters we have considered in this study, we find that there are 15 CaS gene signature in and a 48 CaR gene signature that may impact the prognostic parameters (Table 6, Figure 2). Functional analysis of these genes may give more insight into the mechanisms operating in development of CRPC.

**Table 6:**
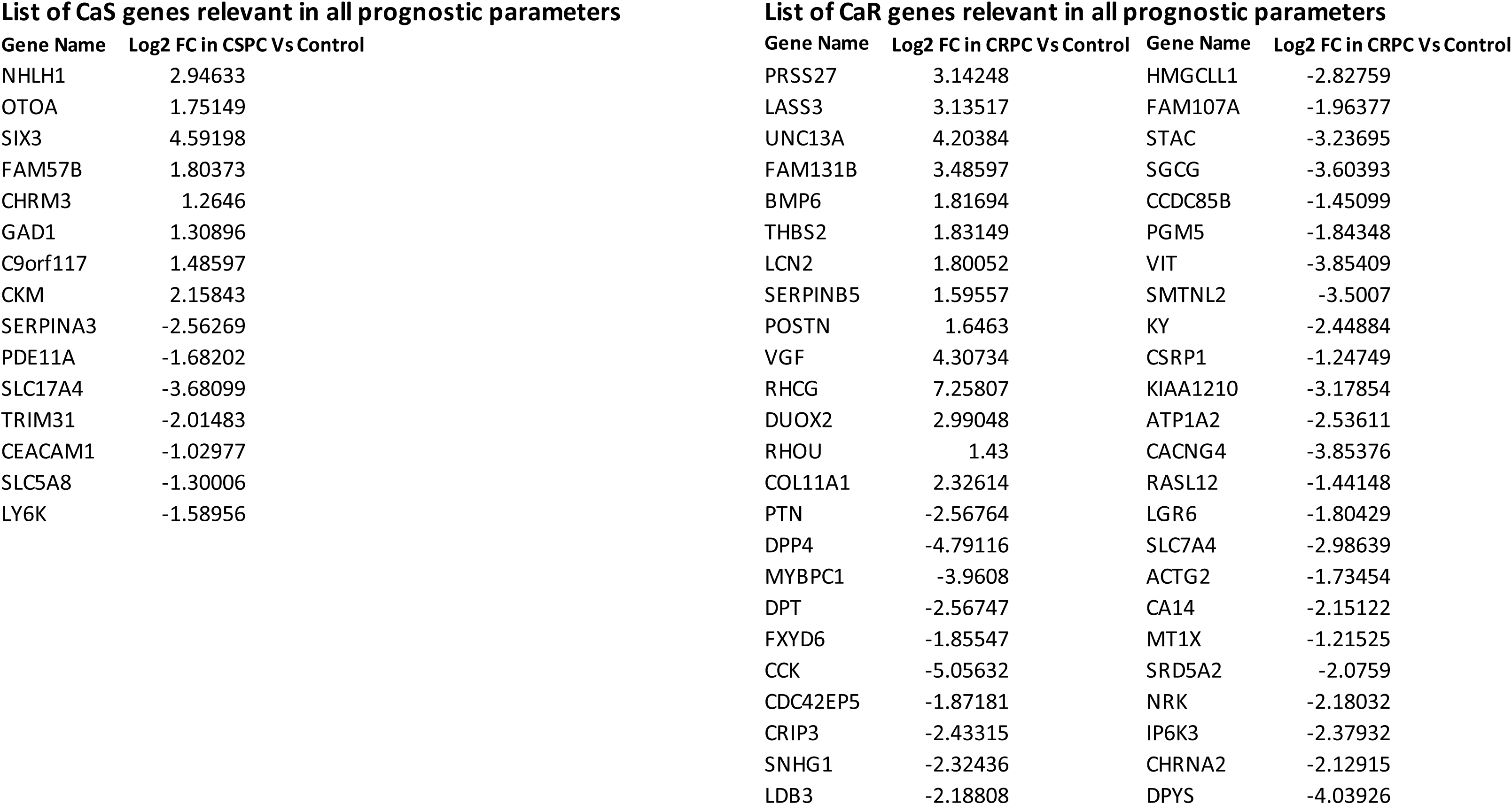
A: List of CaS genes which are relevant in all the parameters of prognosis (15) B: List of CaR genes which are relevant in all the parameters of prognosis (48)

## Declaration

The work described here has been approved by the IEC of both the participating institutions, namely INU and CHG, Bengaluru, India

## Financial Support

The work conducted here has been funded by Rajiv Gandhi University of Health Sciences, Bengaluru. Support by Centre for Human Genetics and Institute of Nephro-Urology is acknowledged.

For part of the duration of the project, PR was supported by Department of Biotechnology, Govt of India under the Ramalingaswami Fellowship.

## Author Contributions

The project was conceived by PR and RK; Sample collection and processing was done by NT and NN respectively; analysis of data has been performed by DJS and SH. All authors have contributed to the manuscript and have approved of it.

DJS, SH and NT have contributed equally for this manuscript.

## Supplementary files

1. Figure 1: Principal Component Analysis of the samples which shows the clustering of the three groups

2. Figure 2: Schematic work flow for the RNA-seq data analysis

Tables

Table 1a: List of differentially expressed genes between Control and Castrate Senstive groups

Table 1b: List of differentially expressed genes between Control and Castrate Resistant groups

Table 1c: List of differentially expressed genes common to both Castrate Senstive and Castrate resistant groups

Table 2: List of genes common to CHG RNA-seq data and TCGA

A. List of CaS genes with fold changes from RNA-seq data

B. List of CaR genes with fold changes from RNA-seq data

## Notes

### Competing Interest Statement

The authors have declared no competing interest.

