## Supplementary figures and images for "Gene expression signature of castrate resistant prostate cancer"

### Supplemantary figure 1

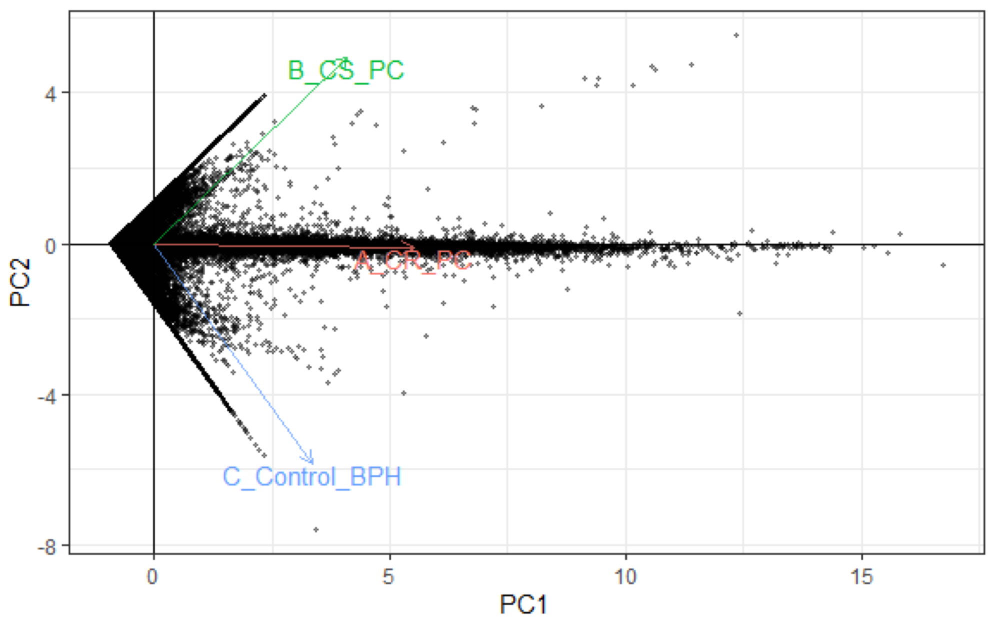


Supplementary Figure 1

### Supplementary figure 2

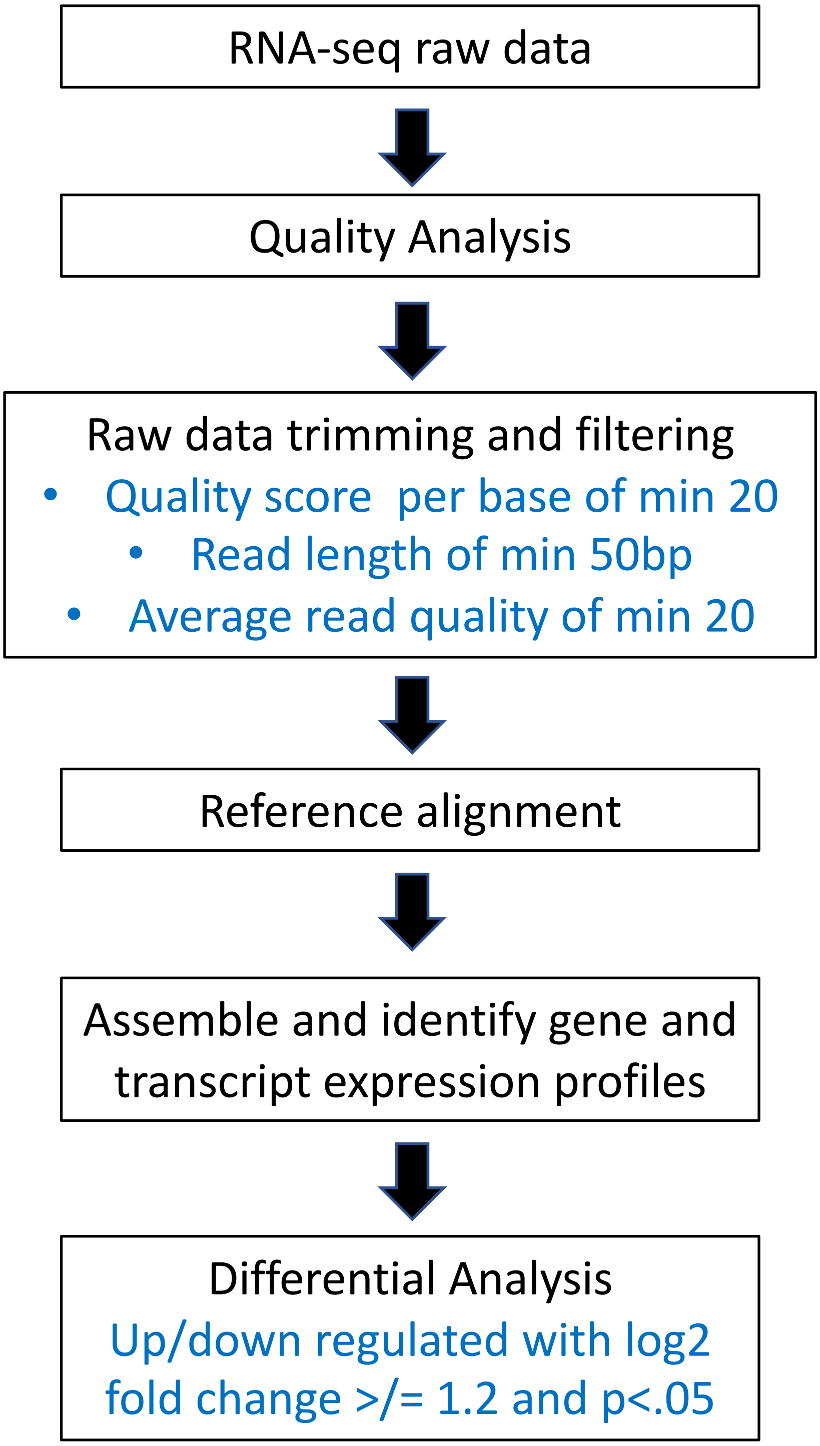


Supplementary Figure 2
